# Understanding implicit sensorimotor adaptation as a process of proprioceptive re-alignment

**DOI:** 10.1101/2021.12.21.473747

**Authors:** Jonathan S. Tsay, Hyosub E. Kim, Adrian M. Haith, Richard B. Ivry

**Author notes:** **Corresponding author Information:** Name: Jonathan Tsay & Richard Ivry, &, Address: 2121 Berkeley Way, Berkeley, CA 94704.

## Abstract

Multiple learning processes contribute to successful goal-directed actions in the face of changing physiological states, biomechanical constraints, and environmental contexts. Amongst these processes, implicit sensorimotor adaptation is of primary importance, ensuring that movements remain well-calibrated and accurate. A large body of work on reaching movements has emphasized how adaptation centers on an iterative process designed to minimize visual errors. The role of proprioception has been largely neglected, thought to play a passive role in which proprioception is affected by the visual error but does not directly contribute to adaptation. Here we present an alternative to this visuo-centric framework, arguing that that implicit adaptation can be understood as minimizing a proprioceptive error, the distance between the perceived hand position and its intended goal. We use this proprioceptive re-alignment model (PReMo) to re-examine many phenomena that have previously been interpreted in terms of learning from visual errors, as well as offer novel accounts for unexplained phenomena. We discuss potential challenges for this new perspective on implicit adaptation and outline a set of predictions for future experimentation.

## I. Implicit adaptation of the sensorimotor system

Motor adaptation is an essential feature of human competence, allowing us to flexibly move in novel and dynamic environments (Kim et al., 2020; Krakauer et al., 2019; Ryan Morehead & de Xivry, 2021; Shadmehr et al., 2010). A sailor adjusts her sails in response to variations in the wind; a basketball player fights against fatigue to maintain a similar force output. Motor adaptation refers to the processes that ensure well-learned movements remain accurate across a broad range of contexts.

Motor adaptation is not a unitary operation but relies on multiple learning processes. Paralleling the memory literature, one broad distinction can be made between processes that are under conscious control and those that operate outside awareness. To continue with the sailing example, a skilled skipper can strategically adjust the sails to achieve a desired heading, while implicitly maintaining that heading based on subtle fluctuations in the rope’s tension. The interplay of explicit and implicit processes in sensorimotor adaptation has been the focus of many studies over the past decade. Whereas the former is volitional and well-suited for rapid modifications in behavior, the latter occurs automatically and operates over a slower time scale (Hegele & Heuer, 2010; Huberdeau et al., 2019; McDougle et al., 2016; Werner et al., 2015).

Computationally, explicit and implicit processes for adaptation are constrained to solve different problems: Whereas explicit processes focus on goal attainment, implicit processes are designed to ensure that the selected movement is flawlessly executed (Taylor & Ivry, 2011). Consistent with this distinction, the deployment of aiming strategies to offset an experimentally imposed perturbation requires prefrontal control (Anguera et al., 2010; Benson et al., 2011; Taylor & Ivry, 2014), whereas implicit adaptation is dependent on the integrity of the cerebellum (Butcher et al., 2017; Haar & Donchin, 2020; Hadjiosif et al., 2014; Izawa et al., 2012; Schlerf et al., 2013; Taylor et al., 2010; Tseng et al., 2007; Tzvi et al., 2021).

One paradigmatic way to study motor adaptation is to introduce a visuomotor perturbation between the motion of the arm and the corresponding visual feedback. Historically, such visuomotor perturbations were accomplished with prism glasses that introduced a translation in the visual field (Helmholtz, 1924; Kitazawa et al., 1995; Petitet et al., 2018; Redding & Wallace, 2001). Nowadays, motion tracking and digital displays enable more flexible control over the relationship between hand position and a feedback signal (Krakauer et al., 2005, 2000). In a typical study, participants are instructed to make reaching movements towards a visual target on a horizontally mounted computer monitor (Figure 1A). By positioning the hand below the display, vision of the hand is occluded. However, a visual cursor is presented on the monitor to indicate hand position, a signal that is readily incorporated into the body schema if its spatial and temporal properties are correlated with the movement. After a few reaches to familiarize the participant with the task environment, a rotation (e.g., 45°) is introduced between the motion of the hand and the visual cursor. If participants continued to move directly to the target, the cursor would miss the target, introducing a visual error. Over several reaches, participants adapt to this perturbation, with the hand’s heading angle shifted in the opposite direction of the rotation.

**Figure 1.**
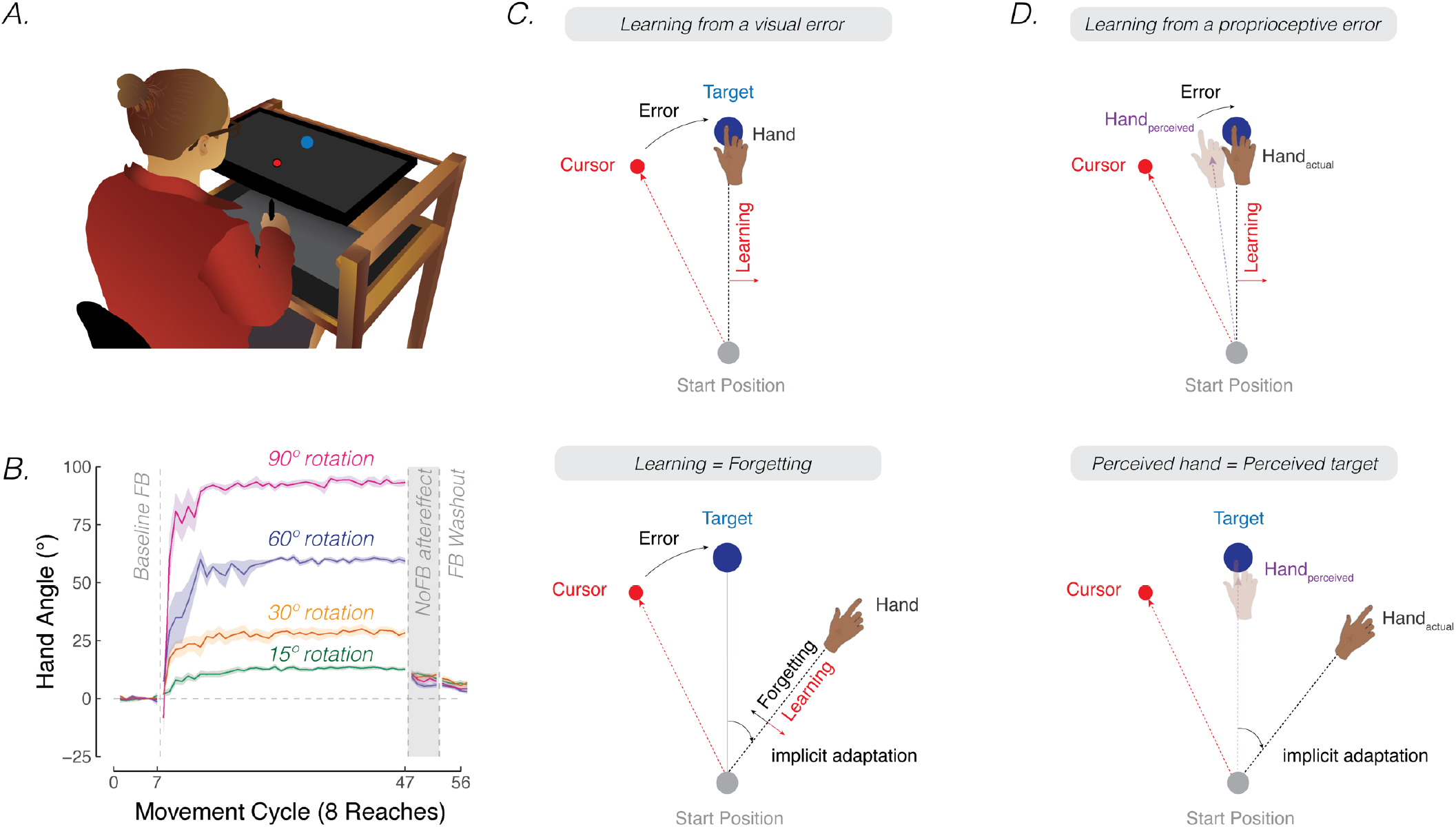
Contrasting visuo-centric and proprioceptive-centric views of implicit motor adaptation. **A)** Experiment setup. **B)** Mean time courses of hand angle for 15° (green), 30° (yellow), 60° (purple), and 90° (pink) rotation conditions from Bond and Taylor (2015). Hand angle is presented relative to the target (0°) during veridical feedback, rotation, and no-feedback trials (grey background). Hand angle is similar during the no-feedback trials for all four perturbation sizes, indicating equivalent implicit adaptation. Shaded region denotes SEM. Note that Bond and Taylor (2015) used eight target locations, and thus, had eight reaches per cycle. **C)** The cursor feedback (red dot) follows a trajectory that is rotated relative to the line connecting the start position and target (blue dot). With visual clamped feedback, the angular trajectory of the visual cursor is not spatially tied to the angular trajectory of the participant’s (hidden) hand but follows a trajectory that is at an invariant angle relative to the target. Despite awareness of the manipulation and instructions to always reach directly to the target, participants show a gradual change in heading direction that eventually reaches an asymptote. According to visuo-centric models, the goal of implicit adaptation is to minimize a visual error (i.e., error = visual cursor – target; upper panel), with the extent of implicit adaptation being the point of equilibrium between learning and forgetting (lower panel). **D)** According to the proprioceptive re-alignment model (PReMo), the goal of implicit adaptation is to minimize a proprioceptive error (i.e., error = perceived hand position – target, upper panel). The perceived (shaded) hand position is influenced by the actual, the expected (based on efference copy), and seen (i.e., visual cursor) hand location. The extent of implicit adaptation corresponds to the point in which the perceived hand position is at the target (lower panel).

The aggregate behavioral change in response to a large perturbation is driven by a combination of strategic aiming and implicit adaptation. One source of evidence originates from a study examining how people respond to visuomotor rotations of varying sizes (Bond & Taylor, 2015). Explicit strategy use, as measured by verbal aim reports, was dominant when the error size was large, producing corrective adjustments in reaching direction that scaled with the size of the rotation (Figure 1B). Yet, the extent of implicit adaptation, as measured by no-feedback aftereffects trials in which participants were instructed to “move directly to the target” without re-aiming, remained constant for perturbations ranging from 15°to 90°. Thus, while explicit re-aiming can flexibly compensate for errors of varying sizes, implicit adaptation saturates, at least for large errors.

The rigidity of implicit adaptation is evident in a variety of other methods (Hegele & Heuer, 2010; Maresch et al., 2020; Mazzoni & Krakauer, 2006; Taylor et al., 2014; Werner et al., 2015). The “visual clamped feedback” task provides an especially striking method to study implicit adaptation without contamination from explicit processes (R. Morehead et al., 2017). With clamped feedback, the angular trajectory of the cursor is invariant with respect to the target, always following a trajectory that is offset from the target by a fixed angle. As such, the direction in which the cursor moves is not contingent on the direction of the participant’s movement. Participants are instructed to always reach directly to the target and ignore the visual cursor. Despite being fully aware of the manipulation, participants adapt, with the heading angle shifting in the opposite direction of the rotation in an automatic and implicit manner. Although the size of the visual error never changes, adaptation eventually reaches an upper bound, averaging between 15° - 25° away from the target. Consistent with the results of Bond and Taylor (Bond & Taylor, 2015), this asymptote does not vary across a wide range of clamped rotation sizes (Kim et al., 2018; Neville & Cressman, 2018; Tsay, Lee, et al., 2021).

## II. The visuo-centric view of implicit sensorimotor adaptation

Implicit adaptation in response to visuomotor perturbations has been framed as an iterative process, designed to minimize a *visual* error (Cheng & Sabes, 2006; Donchin et al., 2003; Herzfeld et al., 2014; Kim et al., 2018; Mazzoni & Krakauer, 2006; R. Morehead et al., 2017; Thoroughman & Shadmehr, 2000; Wolpert et al., 1998). The visual error experienced on the previous trial is used to modify the visuomotor map, such that the motor command on a subsequent trial will be adjusted to reduce that error. According to this visuo-centric view, the extent of implicit adaptation represents a point of equilibrium, one at which the trial-by-trial change in heading angle in response to the visual error is counterbalanced by the trial-by-trial decay (“forgetting”) of this modified visuomotor map back to its baseline, default state (Ryan Morehead & Smith, 2017) (Figure 1C, Appendix).

A visuo-centric perspective on adaptation is appealing. Not only does it fit with a zeitgeist which holds vision as a “dominant” sense, but it also matches our intuition of how we view task success: In day-to-day life, we frequently interact with visual objects, whether it be picking up a glass of water or moving the computer mouse over a desired icon. When a perturbation is introduced, we try to re-establish conditions such that the visual feedback is once again reinforcing. In visuomotor adaptation studies, the experimenter manipulates where the cursor is presented (Krakauer et al., 2000) or when the visual cursor is shown (Brudner et al., 2016; Honda et al., 2012; Kitazawa et al., 1995; Wang et al., 2021). The resultant change in hand trajectory is interpreted as a response to nullify the visual error. A dramatic demonstration of visual dominance comes from the study of deafferented individuals who have lost their sense of proprioception and haptics. Despite their sensory loss, deafferent individuals adapt in a similar manner as those observed in control participants (Bernier et al., 2006; Sarlegna et al., 2010), indicating that vision alone is sufficient to drive implicit adaptation (Blouin et al., 1993; Fleury et al., 1995; Lefumat et al., 2016; Sarlegna et al., 2010; Yousif et al., 2015).

## III. The neglected role of proprioception

Despite its appeal, the visuo-centric view is an oversimplification. The brain exploits all of our senses: While olfaction may not be essential for precisely controlling the limb, proprioception, the perception of body position and body movement, is certainly critical for motor control (Sober & Sabes, 2003, 2005). The classic work of Mott and Sherrington at the end of the 19^th^ Century demonstrated that surgical deafferentation of an upper limb produced severe disorders of movement in the monkey (Mott & Sherrington, 1895). The actions of the animal indicated that the intent was intact, but the movements themselves were clumsy, inaccurate, and poorly coordinated (Bossom, 1974; Munk, 1909). Humans who suffer neurological disorders resulting in deafferentation show a surprising capability to produce well-practiced movements, yet these individuals have marked deficits in feedback control (Rothwell et al., 1982; Sanes et al., 1985). Indeed, recent work indicates that healthy participants rely almost exclusively on proprioceptive information for rapid feedback control, even when visual information about the limb is available (Crevecoeur et al., 2016).

A large body of work also underscores the important role of proprioception in implicit adaptation. First, deafferented individuals fail to generate specific patterns of isometric and isotonic muscle contractions in a feedforward manner as they initiate rapid elbow flexion (Forget & Lamarre, 1987; Gordon et al., 1995). Second, neurologically healthy and congenitally blind individuals can adapt to a force-field perturbation without the aid of vision, presumably relying solely on proprioceptive input (DiZio & Lackner, 2000; Franklin et al., 2007; Marko et al., 2012; Striemer et al., 2019). Third, when opposing visual and proprioceptive errors are provided, aftereffects measured during the no-feedback block after adaptation are in the direction counteracting the proprioceptive error instead of the visual error. That is, implicit adaptation seems to give proprioceptive errors higher priority compared to visual errors (Hayashi et al., 2020) (also see: (Haswell et al., 2009)).

Despite these observations, the computational role of proprioception in implicit adaptation is unclear. In some models, proprioception is seen as playing a passive role, a signal that is biased by vision but does not drive implicit adaptation (Mattar et al., 2013; Ohashi, Gribble, et al., 2019; Ohashi, Valle-Mena, et al., 2019). Other models consider a contribution of proprioception to implicit adaptation, but the computational principles of how this information is used have not been elucidated (Rossi et al., 2021; Ruttle et al., 2021; Salomonczyk et al., 2013; Zbib et al., 2016).

In this review article, we present a new model of sensorimotor adaptation, the proprioceptive re-alignment model (PReMo). The central premise of the model is that proprioceptive error is the primary driver of implicit adaptation, solving the computational problem of ensuring an alignment of the perceived and desired position of the hand. After laying out a set of core principles motivating the model, we present a review of the adaptation literature through this new lens to offer a parsimonious and novel account of a wide range of phenomena. Given that much of the evidence we review is largely based on correlational studies, we conclude by outlining directions for future experimental manipulations that should provide strong tests of PReMo.

## IV. Interaction of Visual and Proprioceptive Information

Perception depends on a combination of multisensory inputs, contextualized by our expectations (Rock, 1983). In reaching to pick up objects in the environment, the location of the hand is specified by afferent inputs from muscle spindles that convey information about muscle length/velocity as well as by visual information relayed by photoreceptors in the eye (Proske & Gandevia, 2012). Estimates of hand position from these signals, however, may not be in alignment due to noise in our sensory systems or perturbations in the environment. To resolve such disparities, the brain shifts the perception of discrepant representations towards one another – a phenomenon known as sensory recalibration.

In the case of a visuomotor rotation, exposure to the systematic discrepancy between vision and proprioception results in a reciprocal interaction between the two sensory signals: The seen (visual) hand location shifts towards the felt (proprioceptive) hand location (i.e., visual shift) (Rand & Heuer, 2019a; Simani et al., 2007) and the felt hand location shifts towards the seen hand location (i.e., proprioceptive shift) (Burge et al., 2010; Cressman & Henriques, 2010; Recanzone, 1998; Synofzik et al., 2008, 2006; K. van der Kooij et al., 2013; Katinka van der Kooij et al., 2016). However, at least for large discrepancies, this visuo-proprioceptive recalibration does not result in a unified percept. Rather the shift within each modality saturates as the visuo-proprioceptive discrepancy increases. For example, visuomotor rotations of either 15° or 30° will result in a 5° shift in proprioception towards the visual cursor and a 1° shift in vision towards the actual hand position (Rand & Heuer, 2019a; Salomonczyk et al., 2013; Simani et al., 2007). [Footnote 1: One critical difference between sensory recalibration and sensory integration is in terms of the resulting percept. As commonly conceptualized (but see Footnote 2), sensory integration results is a unified percept of hand position by combining sensory information in a weighted fashion based on their relative uncertainties (Burge et al., 2008; Van Beers et al., 1999). It is a transient phenomenon that is measured only when *both* modalities are present. In contrast, sensory recalibration is an enduring bias that can be observed when each sensory modality is assessed independently.]

Sensory expectations also play a role in sensory recalibration (‘t Hart & Henriques, 2016). For instance, perception of the moving limb is biased towards the direction of the motor command (e.g., a visual target) (Bhanpuri et al., 2013; Blakemore et al., 1998; Gaffin-Cahn et al., 2019; Gandevia & McCloskey, 1978; Kilteni et al., 2020; Lanillos et al., 2020; McCloskey et al., 1974). One model suggests that the cerebellum receives an efference copy of the descending motor command and generates a prediction of the expected sensory consequences of the movement. This prediction is widely relayed to different regions of the brain, providing a form of predictive control (Grüsser, 1994; Sperry, 1950; von Holst & Mittelstaedt, 1950; Wolpert & Miall, 1996). In sum, sensory recalibration seeks to form a unified percept of hand position by combining sensory inputs and sensory expectations (Körding & Wolpert, 2004).

Sensory recalibration has several notable features: First, sensory recalibration effects are enduring and can be observed even when each sensory modality is assessed alone. For instance, after exposure to a visual perturbation, a visual shift is observed when participants are asked to judge the position of a briefly flashed visual cursor (Simani et al., 2007). Similarly, a proprioceptive shift is observed when participants locate their unseen hand using a touch screen (Ruttle et al., 2021). Second, these shifts occur rapidly (Ruttle et al., 2016, 2021, 2018), with both the visual and proprioceptive shift reaching asymptotic values within just a few reaches after the introduction of a visuomotor perturbation (Ruttle et al., 2016). In the following section, we formalize how sensory recalibration during visuomotor adaptation drives implicit adaptation. Third, as noted above, while the extent of recalibration is a fixed ratio for small visuo-proprioceptive discrepancies (Zaidel et al., 2011), the magnitude of the change within each modality exhibits marked saturation (Salomonczyk et al., 2013; Tsay, Kim, Parvin, et al., 2021; Tsay, Parvin, et al., 2020).

## V. The proprioceptive re-alignment model (PReMo)

Reaching movements are enacted to transport the hand to an intended goal. In most situations, that goal is to pick up an object such as the fork at the dinner table. The resultant feedback allows the brain to evaluate whether the movement ought to be modified. This feedback can come from vision, seeing the hand miss the fork, as well as proprioception, gauging the position of the hand as it misses the fork. The sensorimotor system exploits these multiple cues to build a unified percept of the position of the hand. When the action falls short of meeting the goal – the fork is missed or improperly grasped – adaptation uses an error signal to recalibrate the system. In contrast to visuo-centric models, we propose that the fundamental error signal driving adaptation is proprioceptive, the mismatch between the perceived and desired hand position (Figure 1D, top). From this perspective, the upper bound of implicit adaptation will correspond to the point at which the hand is perceived to be aligned with the target. In this section, we formally develop this proprioceptive re-alignment model (PReMo).

### Perceived hand position is determined by sensory recalibration

As noted above, perceived hand position is determined by a multitude of sensory inputs and sensory expectations. Prior to the crossmodal interaction between vision and proprioception, we assume that the system generates an optimal *intramodal* estimate of hand position using the weighted average of the actual position of the hand (*x*_*p,t*_) and the expected position of the hand based on an outgoing motor command (Figure 2A-B). This motor command is selected to achieve a proprioceptive goal, *G*_*t*_. Therefore, the proprioceptive *integrated hand position* 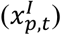 is given by: [Footnote 2: As noted in Footnote 1, the term “integration” is sometimes used to refer to the combination of information from two different sensory modalities (e.g., the *crossmodal* combination of the observed cursor and felt hand position). In this review, we will reserve the term “integration” in an intramodal sense, referring to the combination of the input from a sensory modality and the expected position of that sense based on the outgoing motor command (e.g., for proprioception, the actual and expected hand position; for vision, the actual and expected cursor position). We also note that the proprioceptive movement goal is typically assumed to be the visual target. However, if participants were to use an aiming strategy to compensate for a perturbation, the movement goal would then correspond to the aiming location].

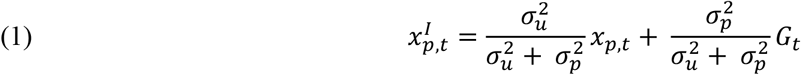

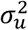 represents the uncertainty of sensory expectations/predictions given a motor command to the goal, which may be influenced by both extrinsic sources of variability (e.g., greater perturbation variability in the environment) and intrinsic sources of variability (e.g., greater motor noise). 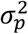 represents uncertainty in the proprioceptive system.

**Figure 2.**
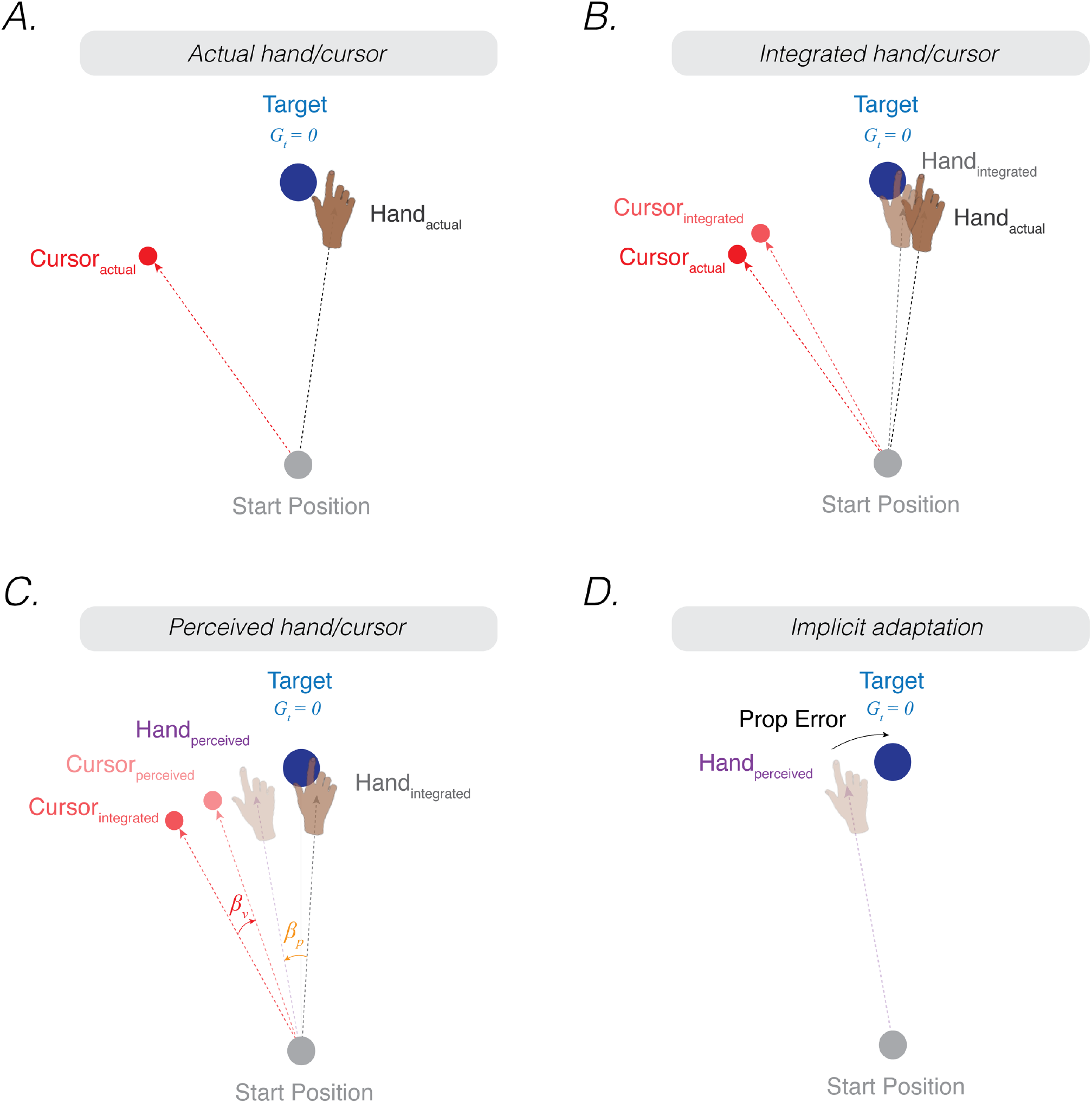
The proprioceptive re-alignment model (PReMo). **A)** When the feedback is rotated, the position of the feedback cursor (red dot) is rotated counterclockwise with respect to the location of the unseen hand (deviation depicted here arises from motor noise). **B)** Due to intramodal integration of sensory input and sensory expectations from the motor command, the integrated hand would lie between the visual target and the actual position of the hand, and the integrated cursor would lie between the visual target and the actual cursor. **C)** Due to crossmodal sensory recalibration, the integrated hand shifts towards the integrated cursor (proprioceptive shift, *β*_*v*_) and the integrated cursor shifts towards the integrated hand (visual shift, *β*_*v*_), forming the perceived hand and perceived cursor locations. **D)** The proprioceptive error (mismatch between the perceived hand position and the target, *G*_*t*_) drives implicit adaptation in the clockwise direction, opposite to the imposed counterclockwise rotation.

Correspondingly, the optimal *intramodal* integrated estimate of the visual cursor position 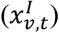 is the weighted average of the actual position of the cursor (*x*_*v,t*_) and the expected position of the cursor based on outgoing motor commands (*G*_*t*_):

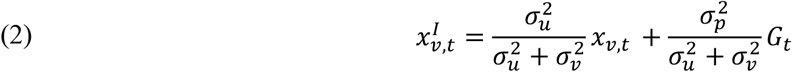

where 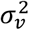 represents uncertainty in the visual system. This intramodal integrated estimate of hand position is recalibrated crossmodally by vision (proprioceptive shift, (*β*_*t*_), resulting in a perceived hand position 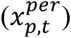 (Figure 2C):

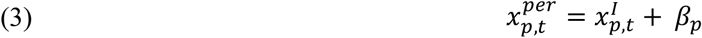

Correspondingly, the intramodal integrated estimate of cursor position is recalibrated crossmodally by proprioception (visual shifts, (*β*_*v*_), resulting in a perceived cursor position 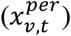:

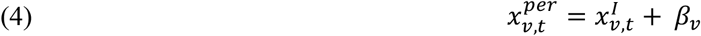

While the exact computational rules that govern the magnitude of crossmodal shifts (*β*_*p*_, *β*_*v*_)remain an active area of research (Hong et al., 2020), we assume that the perceptual shifts follow three general principles based on observations reported in the previous literature:

A. For small discrepancies, the degree of crossmodal recalibration is a fixed ratio (i.e., *η*_*p*_, *η*_*v*_) of the visuo-proprioceptive discrepancy (i.e., 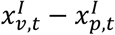). For larger discrepancies, the magnitude of the shift for each modality saturates (i.e., *β*_*p,sat*_, *β*_*v,sat*_)(Ruttle et al., 2021; Salomonczyk et al., 2013; Synofzik et al., 2008; ‘t Hart et al., 2020):

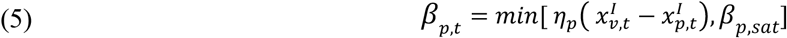

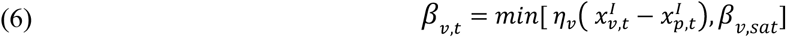

B. Visual and proprioceptive shifts are global (Rand & Heuer, 2019b, 2020; Simani et al., 2007). This implies that the perceived hand position is not only shifted in the region of space near the biasing source (i.e., the target or feedback cursor), but will also be shifted in the same direction across the workspace (e.g., at the start position).

C. Crossmodal recalibration decays in the absence of visual feedback (the rate of decay: *A*) (Babu et al., 2021).

### Proprioceptive error signal drives implicit adaptation

As stated in the previous section, the motor system seeks to align the perceived hand position with the movement goal. A proprioceptive shift induced by a visuo-proprioceptive discrepancy will mis-align the perceived hand position with the movement goal, resulting in a proprioceptive error:

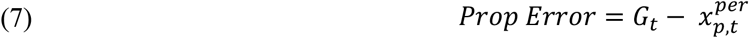

The proprioceptive error is used to update the sensorimotor map such that a subsequent motor command will bring the hand position closer to being in alignment with the target. As with most state space models, this update process operates with a learning rate (*K*):

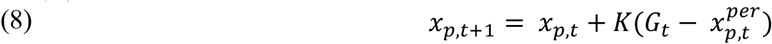

A key assumption of PReMo is that the upper bound of adaptation 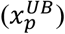 is determined as the position at which the proprioceptive error is eliminated (i.e., the perceived hand position = the perceived motor goal). By rearranging the terms in Eq 1 and 3, and assuming that the movement goal is at the target (i.e., 0°), the upper bound of adaptation is given by:

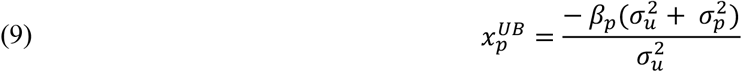

This equation has several important implications for the upper bound of implicit adaptation. First, the upper bound of adaptation will increase with proprioceptive uncertainty 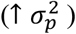 (Eq 9) and the size of the proprioceptive shift (↑ *β*_p_) (Eq 9). Second, rewriting the equation shows that the upper bound of adaptation will be attenuated when there is an increase in the noise associated with sensory expectations of the motor command 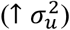 (Eq 10) (see Supplemental section titled: *Proprioceptive shift does not correlate with proprioceptive variability*):

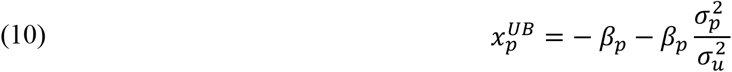

Third, assuming that the proprioceptive shift saturates (*β*_*p,sat*_) for a wide range of visuo-proprioceptive discrepancies, the proprioceptive error will saturate (Eq 11). As such, trial-by-trial motor updates (*U*_*sat*_, Eq 12) and the extent of implicit adaptation (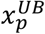,Eq 13) will also saturate:

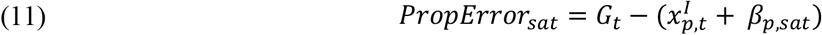

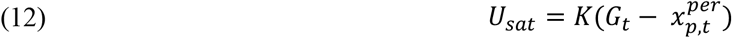

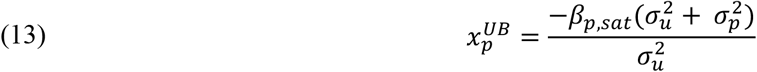

Finally, and perhaps most strikingly, the perceived location of the hand will follow a non-monotonic pattern. There will be an initial bias away from the target towards the visual cursor due to the proprioceptive shift (crossmodal recalibration between vision and proprioception). The resulting proprioceptive error would drive the hand in the opposite direction, away from the visual cursor. Thus, the perceived hand position would gradually rebound back to the target until the proprioceptive error is nullified. This last prediction encapsulates the essence of PReMo.

## VI. Empirical support for the proprioceptive re-alignment model

In this section, we evaluate the evidence in support of PReMo, focusing on the predictions outlined above.

### Feature 1. Implicit adaptation is correlated with proprioceptive shift

A core observation that spurred the development of PReMo is the intimate link between the proprioceptive shift and extent of implicit adaptation. One common method to quantify measures of proprioception involves asking participants to report the position of their hand after passive displacement (Figure 3A). The psychometric function derived from these reports is used to estimate the participants’ bias and variability (see Feature 3 below for an extended discussion on proprioceptive variability). The proprioceptive judgements (i.e., “indicate where you feel your hand”) are usually obtained before and after the visual feedback is perturbed, and as such, can be used to quantify the proprioceptive shift (i.e., change in proprioceptive bias, (*β*_*p*_ note that this measure is not the same as perceived hand position, 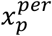). Across a range of experiments, two notable characteristics stand out: 1) The proprioceptive shift saturates at ∼5º and 2) reaches an asymptotic value after only a couple of trials of exposure to a visuo-proprioceptive discrepancy (Block & Bastian, 2011; Cressman & Henriques, 2010; Gastrock et al., 2020; Modchalingam et al., 2019; Rand & Heuer, 2019a).

**Figure 3.**
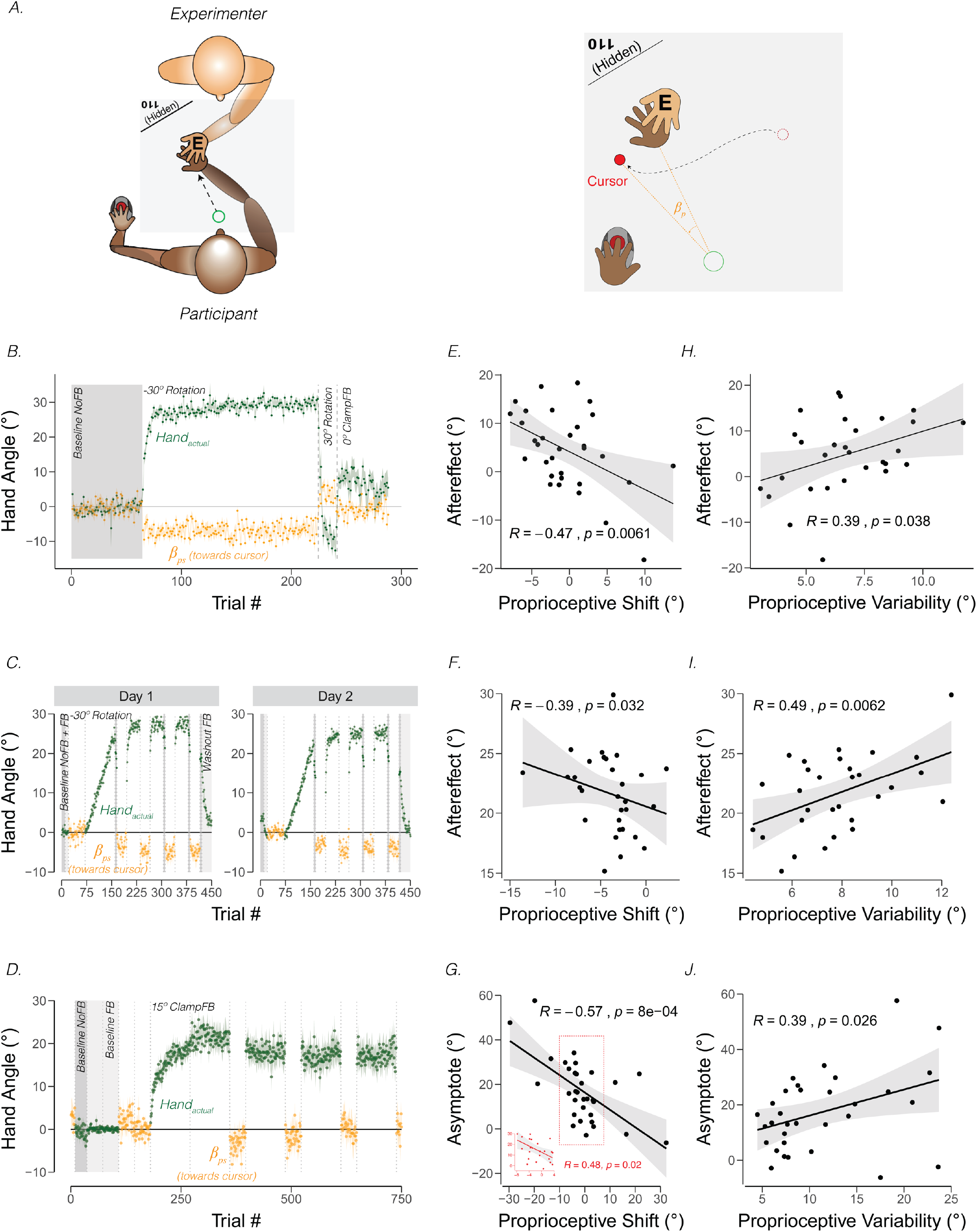
Proprioceptive shift and variability correlate with the upper bound of adaptation. **A)** Experimental setup for proprioceptive probe trials in Tsay et al (2021). The experimenter sat opposite the participant and moved the participant’s hand from the start position to a location specified in the corner of the monitor (e.g., 110º) that was only visible to the experimenter. After the participant’s hand was passively moved to the probe location, a cursor appeared at a random position on the screen (right panel). The participant used their left hand to move the cursor to the perceived hand position. A similar method was used in Ruttle et al 2021), but instead of the experimenter, a robot manipulandum was programmed to passively move the participant’s arm. **B)** Abrupt visuomotor rotation design from Ruttle et al (2021, Exp 1). After a baseline veridical feedback block, participants were exposed to a -30° rotated cursor feedback block, a 30° rotated feedback block, and a 0° clamped feedback block. Vertical dotted lines indicate block breaks. Green dots denote hand angle. Orange dots denote proprioceptive probe trials. Shaded error bars denote SEM. **C)** Gradual visuomotor rotation design from Tsay et al (2021, Exp 1). After baseline trials without feedback (dark grey) and veridical feedback (light grey), participants were exposed to a perturbation that gradually increased to -30º and then held constant. There were periodic proprioceptive probe blocks (orange dots) and no feedback motor aftereffect blocks (dark grey). **D)** Clamped rotation design from Tsay et al (2021, Exp 2). After a period of baseline trials, participants were exposed to clamped visual feedback that moves 15° away from the target. **E – G)** Correlation between proprioceptive shift and the extent of implicit adaptation. Note that the correlations are negative because a leftward shift in proprioception (towards the cursor) will push adaptation further to the right (away from the target and in the opposite direction of the cursor). Black dots represent individual participants. **H – J)** Correlation between variability on the proprioceptive probe trials during baseline and the extent of implicit adaptation in the three experiments depicted in B-D.

[Footnote 3: Proprioceptive recalibration may differ between experimental setups in which the hand movement and visual feedback are co-planar or occur in different planes (e.g., horizontal hand movement with visual feedback on a vertically aligned monitor). In the latter case the proprioceptive estimate requires an extra coordinate transformation. Nevertheless, PReMo can account for proprioceptive recalibration/shifts if provided with a representation of the actual hand position, predicted hand position, and visual feedback regarding hand position, with the orthogonal case requiring a coordinate transformation. There is considerable behavioral and neural evidence showing that we perform coordinate transformations with considerable flexibility (Miller et al., 2018). Indeed, this ability allows us to endow prosthetics and tools with “proprioception” (Kieliba et al., 2021), perceiving them as extensions of our own bodies.]

Across individuals, the magnitude of the proprioceptive shift can be correlated with the extent of adaptation, operationalized as the magnitude of the aftereffect obtained after exposure to a visuomotor rotation (Ruttle et al., 2021; Tsay, Kim, Parvin, et al., 2021) or as the asymptotic change in reaching angle following exposure to visual clamped feedback (Tsay, Kim, Parvin, et al., 2021). As can be seen in Figure 3 (Panels E-G), the magnitude of the shift is negatively correlated with the upper bound of implicit adaptation. That is, the more proprioception shifts towards the cursor position, the greater the extent of implicit adaptation away from the perturbed cursor. A similar pattern has been observed in many other studies (Clayton et al., 2014; Gastrock et al., 2020; Modchalingam et al., 2019; Salomonczyk et al., 2011, 2013; Simani et al., 2007).

The correlation between proprioceptive shift and the upper bound of adaptation is in accord with PReMo (Eq. 9). A greater shift in perceived hand location towards the perturbed visual feedback would create a greater misalignment between the perceived hand position and the desired hand position (i.e., the perceived location of the target). As such, one would expect that a larger deviation in hand angle would be required to offset this shift. With the focus on the visual error signal, visuo-centric models of implicit adaptation do not consider how the visual perturbation impacts the perceived hand location. Thus, these models do not predict, or rather are moot on the relationship between proprioceptive shift and the upper bound of adaptation.

In the following section, we will explore five unexplained phenomena of implicit adaptation that can be accounted for by observed features of the shift in proprioception.

### Feature 1, Corollary 1: The rate and extent of implicit adaptation saturates

Many studies of sensorimotor adaptation have examined how the system responds to visual errors of varying size. Standard adaptation tasks that use a fixed perturbation and contingent visual feedback are problematic since behavioral changes that reduce the error also increase task success. To avoid this problem, two basic experimental tasks have been employed. First, the visual perturbation can vary in terms of both size and sign on a trial-by-trial basis, with the rate of implicit adaptation quantified as the change in hand trajectory occurring on trial n + 1 as a function of the visual error experienced on trial n (Hayashi et al., 2020; Marko et al., 2012; Wei & Körding, 2009). By varying the sign as well as the size, the mean visual error is held around 0°, minimizing cumulative effects of learning. Second, clamped visual feedback can be used to look at an extended learning function to a constant visual error signal (Kim et al., 2018; R. Morehead et al., 2017). With these data, one can estimate an initial rate of implicit adaptation (e.g., change over the initial trials in response to a clamp), as well as measure the asymptotic value of adaptation. With traditional adaptation tasks, the asymptote of implicit adaptation can only be measured in an aftereffect block (Bond & Taylor, 2015).

A striking result has emerged from this work, namely that the rate and extent of implicit adaptation is only proportional to the size of the error for small errors before saturating across a broad range of larger errors at around 5° (Figure 4) (Hayashi et al., 2020; Kasuga et al., 2013; Kim et al., 2018; Marko et al., 2012; R. Morehead et al., 2017; Tsay, Lee, et al., 2021; Wei & Körding, 2009). As can be seen in Figure 4A, the rate (e.g., trial-to-trial change in hand angle) is relatively invariant in response to visual errors that exceed 10°.

**Figure 4.**
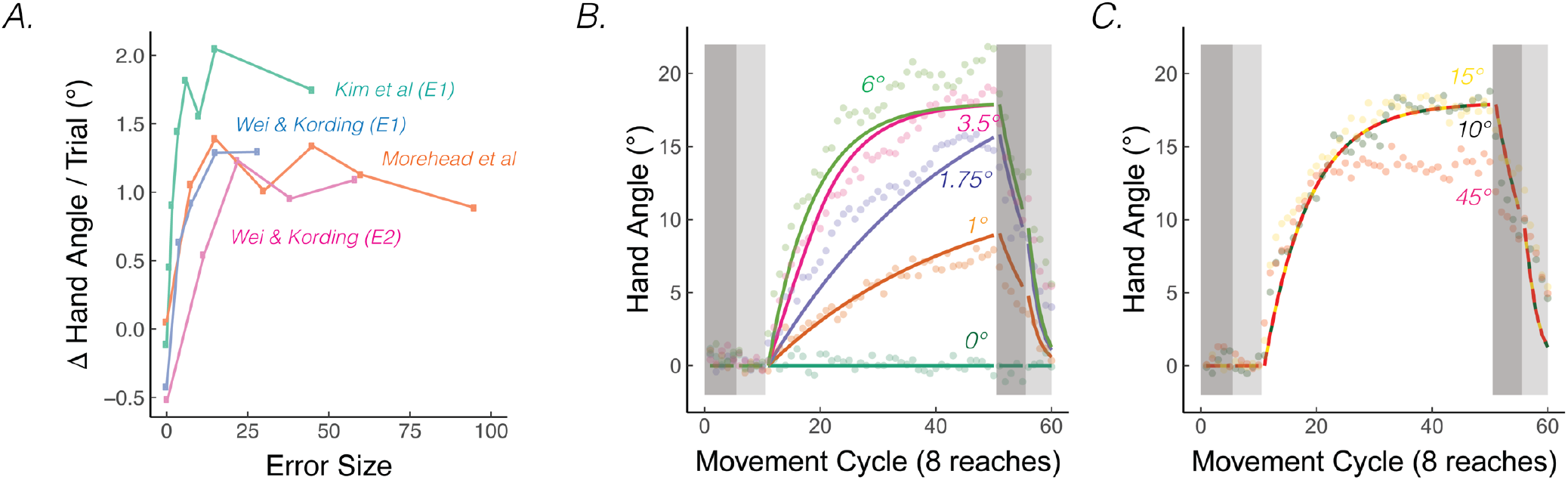
The rate of implicit adaptation saturate. **A)** Early adaptation rates saturate for studies using different methodologies. **B)** Data from Kim et al (2018). Different groups of participants made reaching movements with clamped visual feedback of varying sizes (0° - 45°; groups were divided into two panels for visualization purposes). Groups with smaller clamps (less than 6°) exhibited early adaptation rates that scaled with the size of the clamped feedback, but groups with larger clamped feedback (6° and above) showed a saturated early adaptation rate. Lines denote model fits of the proprioceptive re-alignment model (*R*^2^ = 0.953).

A central feature of the standard visuo-centric model is that the rate and asymptote of implicit adaptation are determined by two fixed parameters, the learning and forgetting rates. By this view, the change in the sensorimotor map following a given trial will be a fixed proportion of the visual error size: That is, rate will scale with error size. Similarly, the asymptote would also scale, reaching a final level at which the change resulting from the response to the visual error on the previous trial is in equilibrium with the amount of forgetting on the previous trial. This visuo-centric perspective cannot explain why the rate and extent of adaptation would saturate.

Various modifications of the visuo-centric model have been developed to accommodate these saturation effects. One such model centers on the idea that the motor system reduces its learning rate in response to large visual errors, an argument that is ecologically grounded in the idea that these large errors arise from rare, external events rather than errors that arise from within the motor system (Shams & Beierholm, 2010; Wei & Körding, 2009). Another model centers on the motor system being limited in its short-term motor plasticity, with the upper bound reflecting the maximum amount of behavioral change the motor system can accommodate (Kim et al., 2018).

PReMo offers a novel account of the saturation effect, one that shifts the focus from the motor system to the sensory system. As noted previously, the size of the proprioceptive shift saturates at a common value (∼5°) across a wide range of visuo-proprioceptive discrepancies (Mostafa et al., 2015; Ruttle et al., 2021; Salomonczyk et al., 2013; Synofzik et al., 2008, 2010, 2006; Tsay, Parvin, et al., 2020). For example, the proprioceptive shift is essentially the same following the introduction of a 15° rotation or a 30° rotation (Tsay, Kim, Parvin, et al., 2021). Since the size of the proprioceptive shift dictates the size of the proprioceptive error, we should expect an invariant rate and extent of motor adaptation in response to visual errors of different sizes (Eq. 11-13). Even under experimental manipulations for which the proprioceptive shift scales with the size of small visual errors, the same scaling is mirrored in the extent of implicit adaptation (‘t Hart et al., 2020), further supporting the link between the proprioceptive shift and implicit adaptation.

### Feature 1, Corollary 2: Proprioceptive shift at the start position explain patterns of generalization

Generalization provides a window into the representational changes that occur during sensorimotor adaptation. In visuomotor rotation tasks, generalization is assessed by exposing participants to the perturbation during movements to a limited region of the workspace and then examining changes in movements made to other regions of the workspace (Ghahramani et al., 1996; Pine et al., 1996). A core finding is that generalization of implicit adaption is local, with changes in trajectory limited to targets located near the training region (Krakauer et al., 2000; Tanaka et al., 2009). These observations have led to models in which generalization is determined by the properties of directionally-tuned motor units, with the extent of generalization dictated by the width of their tuning functions (Tanaka et al., 2009). As such, the error signal that drives implicit adaptation only produces local changes around the location where the error was experienced. From the lens of PReMo, this view of local generalization specifies how implicit adaptation attributable to proprioceptive re-alignment at the training target should affect movements to nearby target locations.

More intriguing, many studies have also found small, but reliable changes in heading direction to targets that are far from the training location, including at the polar opposite direction of training (Krakauer et al., 2000; R. Morehead et al., 2017; Pine et al., 1996; Poh et al., 2021; Taylor et al., 2013). For example, following an exposure phase in which a 45° CCW rotation was imposed on the visual feedback for movements to one target location, a 5° shift in the CW direction was observed for movements to probe locations more than 135° away (Figure 5C, Taylor et al (2013)). These far generalization effects have been hypothesized to reflect some sort of global component of learning, one that might be associated with explicit re-aiming (Hegele & Heuer, 2010; McDougle et al., 2017; McDougle & Taylor, 2019). [Footnote 4: Unlike visuomotor adaptation, force-field adaptation does not appear to produce far generalization (Howard & Franklin, 2015; Rezazadeh & Berniker, 2019). Future research can evaluate how constraints on PReMo vary between different tasks. See Feature 6 about how PReMo generalizes from visuomotor rotation to force-field adaptation]

**Figure 5.**
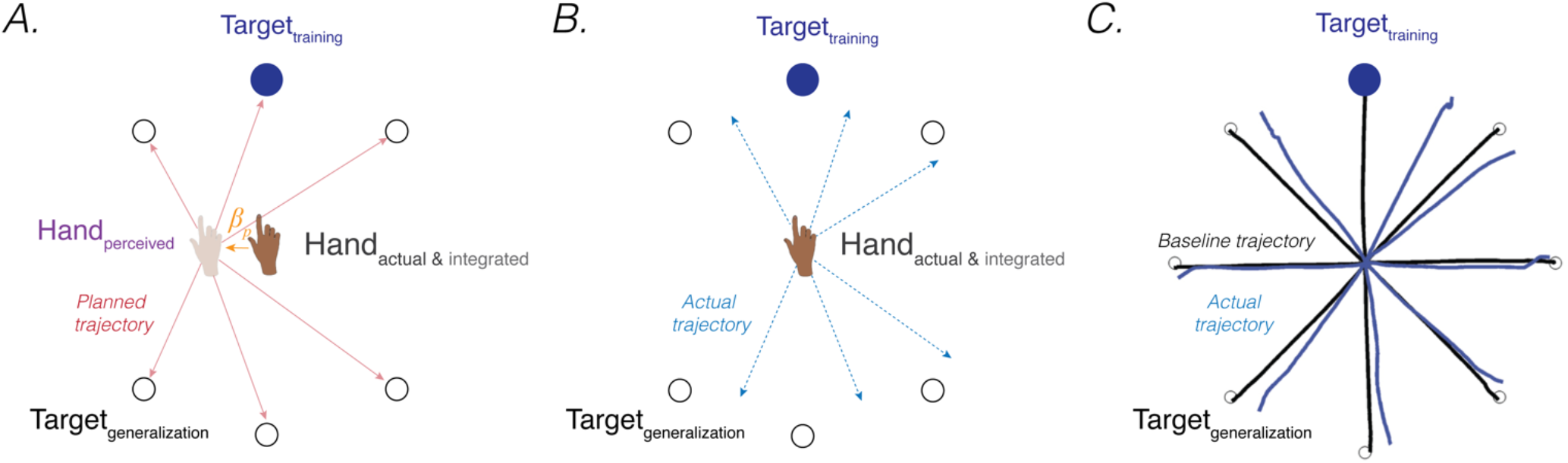
Proprioceptive shift at the start position explain patterns of generalization following a visuomotor rotation. **A)** Planned movement trajectories are formed by participants planning to make a movement initiated at their perceived hand position to the target location (red solid lines). The perceived hand position is assumed to be biased by a proprioceptive shift (*β*_*p*_). **B)** The predicted generalization pattern of the proprioceptive re-alignment model (dashed blue lines; the planned trajectory is initiated from the actual/integrated hand position at the start position). **C)** Pattern of generalization in Taylor et al (2013) assessed during no-feedback generalization trials following training with a 45° CCW rotation in one direction (upward). Black lines are baseline trajectories; blue lines are generalization trajectories. Note match of observed generalization pattern with predicted pattern shown in B), with some trajectories deviated in the clockwise direction, others in the counterclockwise direction, and no change in heading for the reaches along the horizontal meridian.

However, PReMo suggests an alternative interpretation, positing that far generalization arises as a consequence of the cross-sensory recalibration that comes about from exposure to the perturbation. Cross-sensory recalibration has been shown to result in visual and proprioceptive shifts that extend across the training space. That is, when proprioceptive judgements are obtained pre- and post-training, the resulting distortions of vision and proprioception are remarkably similar at the trained and probed locations around the workspace (Cressman & Henriques, 2010; Simani et al., 2007; ‘t Hart & Henriques, 2016; ‘t Hart et al., 2020). Specifically, Simani et al. (2007) observed a robust proprioceptive shift at a generalization target 45° from the training location; this finding was extended in studies from the Henriques’ group, revealing a robust proprioceptive shift as far as 100° from the training target (Mostafa et al., 2015; ‘t Hart & Henriques, 2016; ‘t Hart et al., 2020).

Assuming that the proprioceptive shift induced by a visuomotor rotation also affects the perceived hand location at the start position (an assumption yet to be directly tested), all movement trajectories planned from the perceived (shifted) hand position at the start position to a visual target located at any position within the workspace will be impacted (Sober & Sabes, 2003; Vindras et al., 1998) (Figure 5A). This will yield a pattern of generalization that extends to far probe locations (Figure 5B). Specifically, the planned vector (solid red line; Figure 5A) would result in an actual clockwise movement with respect to the upward trained target (dotted blue line; Figure 5B) but a counterclockwise movement with respect to the bottom generalization targets. PReMo therefore provides a qualitative account of generalization as a combination of 1) a local pattern of generalization of implicit adaptation around the training target caused indirectly by the proprioceptive shift at the target location, and 2) a global pattern of generalization attributable to the proprioceptive shift at the start location. Future experiments should be conducted to quantify the relative contribution of these two components to the global pattern of generalization following adaptation to a visuomotor rotation.

### Feature 1, Corollary 3: Implicit adaptation is correlated with the proprioceptive shift induced by passive movements

Perturbed feedback during passive limb movement can also drive implicit adaptation. A striking demonstration of this comes from a study by Salomonczyk et al. (2013). In the exposure phase (Figure 6A), the participant’s arm was passively moved along a constrained pathway by a robotic device while a cursor moved to a remembered target location (i.e., the target disappeared when the robot-controlled movement started). Across trials, the passive movement of the arm was gradually rotated away from the cursor pathway over trials, eliciting an increasingly large discrepancy between the feedback cursor and perceived motion of the hand. When asked to report their hand position, the participants showed a proprioceptive shift of around 5° towards the visual cursor, comparable to that observed following active movements with perturbed visual feedback in a standard visuomotor rotation paradigm. After the passive perturbation phase, participants were instructed to actively reach to the visual target. These movements showed a motor aftereffect, deviating in the direction opposite to the cursor rotation. Moreover, the size of the aftereffect was correlated with the magnitude of the proprioceptive shift (Figure 6B). That is, participants who showed a greater proprioceptive shift towards the visual cursor also showed a stronger motor aftereffect.

**Figure 6.**
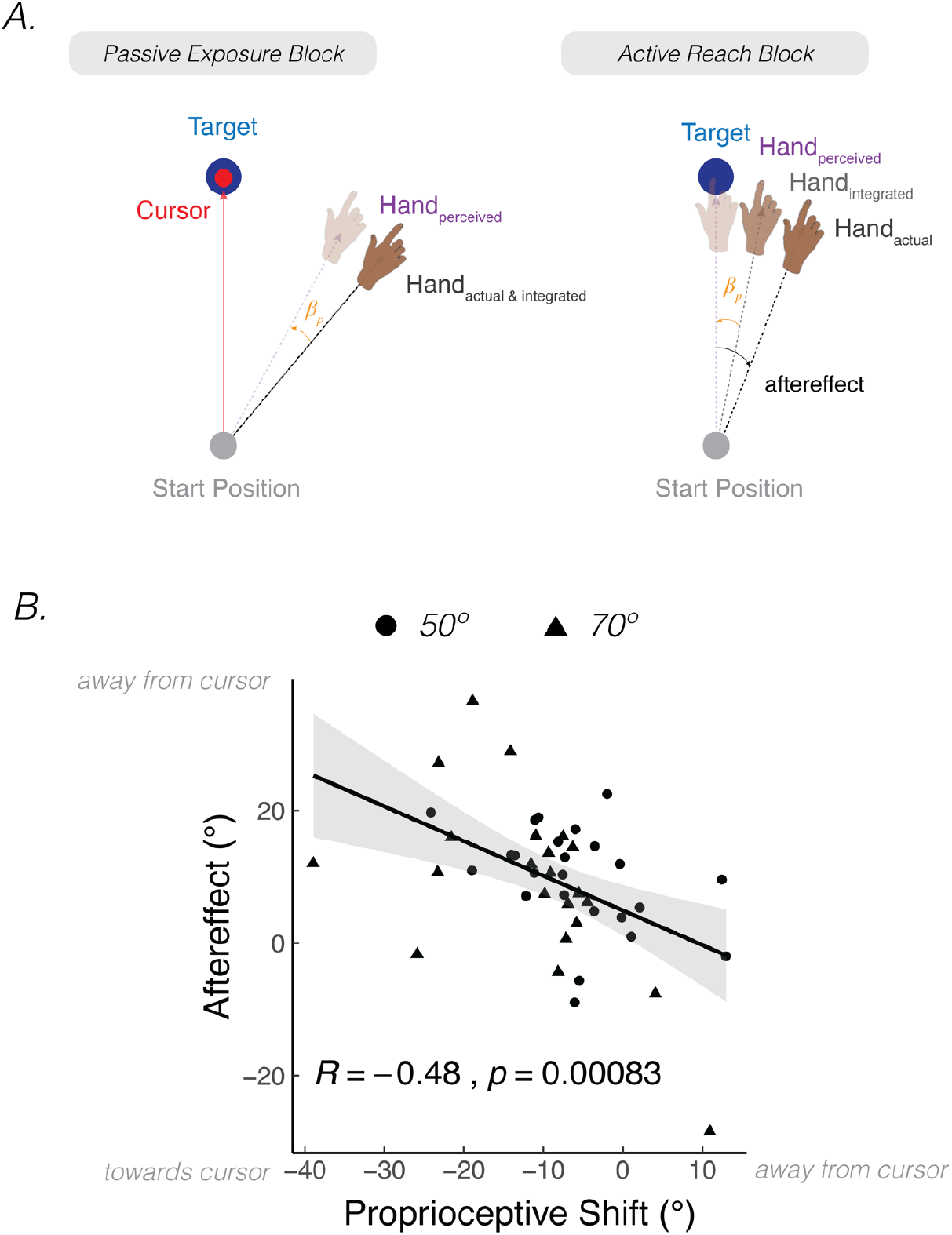
Aftereffects are elicited after passive exposure to a visuo-proprioceptive discrepancy. **A)** In Salomonczyk et al (2013), the hand was passively moved by a robot during the exposure block, with a gradual perturbation introduced that eventually reached 50° or 70° from the target (between-participant design). Simultaneous online visual feedback was provided, with the cursor moving directly to the target position. An aftereffect was measured during the active reach block in which the participant was instructed to reach directly to the target without visual feedback. **B)** Magnitude of shift in perceived hand position assessed during the passive exposure block was correlated with the motor aftereffect (Dots = 50° group; Triangles = 70° group). The more negative values on x-axis indicate a larger proprioceptive shift towards the target. The larger values on the y-axis indicate a larger motor aftereffect.

PReMo can also account for the correlation between a passively-induced proprioceptive shift and the magnitude of implicit adaptation. The proprioceptive shift arises from a discrepancy between the *integrated* position of the visual cursor (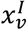; the integration of the actual cursor position and expectation due to the motor command; Eq 2) and the *integrated* position of the hand (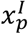; the integration of the actual hand position and the expectation from the motor command; Eq 1). When the hand is passively moved towards a predetermined location, sensory expectations from a motor command are absent. Therefore, the *integrated* positions of the cursor and hand correspond to their *actual* positions. As the integrated (actual) hand is passively and gradually rotated away from the cursor feedback (clamped at the target location), a discrepancy is introduced between the integrated positions. This visuo-proprioceptive discrepancy results in a proprioceptive shift of the integrated hand towards the integrated cursor position, and vice versa.

In contrast, visuo-centric models do not provide an account of how the sensorimotor system would be recalibrated in the absence of movement. Moreover, in this context, adaptation would not be expected given that the visual error was zero (i.e., the cursor always moved directly to the target).

### Feature 1, Corollary 4: Proprioceptive shift and implicit adaptation are both attenuated when visual feedback is delayed

Timing imposes a powerful constraint on implicit adaptation: Delaying the visual feedback by as little as 50-100 ms can markedly reduce the rate of adaptation (Held & Durlach, 1992; Held et al., 1966; Kitazawa et al., 1995). Indeed, evidence of implicit adaptation may be negligible if the visual feedback is delayed by more than 2s (Brudner et al., 2016; Kitazawa et al., 1995). The attenuating effect of delayed visual feedback has been attributed to temporal constraints associated with cerebellar-dependent implicit adaptation. Specifically, while the cerebellum generates sensory predictions with exquisite resolution, the temporal extent of this predictive capability is time-limited, perhaps reflecting the kind of temporal delays that would be relevant for a system designed to keep the sensorimotor system calibrated (Keele & Ivry, 1990; Miall et al., 2007; Wolpert & Miall, 1996; Wolpert et al., 1998). Delaying the visual feedback would presumably result in a weaker sensory prediction error, either because of a misalignment in time between the predicted and actual sensory feedback or because the sensory prediction fades over time. The consequence of this delay would be attenuated implicit adaptation.

Although we have not included temporal constraints in PReMo, it has been shown that delayed visual feedback also attenuates the proprioceptive shift; as such, the model would predict reduced adaptation since the signal driving adaptation is smaller (Eq 1, 3, 5). In visuomotor adaptation studies, the proprioceptive shift is reduced by ∼30% for participants for whom the visual feedback on reaching trials was delayed by 750 ms, relative to those for whom the feedback was not delayed (Debats & Heuer, 2020b; Debats et al., 2021). A similar phenomenon is seen in a completely different task used to study proprioceptive shift, the rubber hand illusion (Botvinick & Cohen, 1998; Longo et al., 2008; Makin et al., 2008). Here timing is manipulated by varying the phase relationship between seeing a brush move along the rubber hand and feeling the brush against one’s own arm: When the two sources of feedback are out of phase, participants not only report less ownership of the rubber hand (indexed by subjective reports), but also exhibit a smaller shift in their perceived hand position towards the rubber hand (Rohde et al., 2011; Shimada et al., 2009).

While studies on the effect of delayed feedback hint at a relationship between proprioceptive shift and implicit adaptation, the supporting evidence is indirect and based on inferences made across several different studies. Future research is required to directly test whether the temporal constraints known to impact implicit adaptation also apply to proprioceptive shift.

### Feature 1, Corollary 5: Proprioceptive shift and implicit adaptation are attenuated by awareness of the visual perturbation

Although we have emphasized that adaptation is an implicit process, one that automatically occurs when the perceived hand position is not aligned with the desired hand position, there are reports that this process is attenuated when participants are aware of the visual perturbation. For example, Neville and Cressman (2018) found that participants exhibited less implicit adaptation (indexed by the motor aftereffect) when they were fully informed about the nature of visuomotor rotation and the strategy required to offset the rotation (see also, (Benson et al., 2011); but see, (Werner et al., 2015)). Relative to participants who were uninformed about the perturbation, the extent of implicit adaptation was reduced by ∼33%. This attenuation has been attributed to plan-based generalization whereby the locus of implicit adaptation is centered on the aiming location and not the target location (Day et al., 2016; McDougle et al., 2017; Schween et al., 2018). By this view, the attenuation is an artifact: It is only reduced when probed at the original target location, since this position is distant from the center of the generalization function. Alternatively, if adaptation and aiming are seen as competitive processes, any increase in strategic re-aiming would be expected to damp down the contribution from the adaptation system (Albert et al., 2020).

For the present purposes, we note that none of the preceding accounts refer to a role of proprioception. However, Debats and Heuer (2020a) have shown that awareness of a visuomotor perturbation attenuates the size of the proprioceptive shift (Debats & Heuer, 2020a). The magnitude of this attenuation in response to a wide range of perturbations (0° - 17.5°) was around 30%, a value similar to the degree to which implicit adaptation was attenuated by awareness in the Neville and Cressman study. A quantitative correspondence in the effect of awareness on proprioceptive shift and implicit adaptation is predicted by PReMo (Eq 9).

Taken together, there is some, albeit relatively thin, evidence that the proprioceptive shift and implicit adaptation may be attenuated by awareness. PReMo provides motivation for further research on this question, both to clarify the impact of awareness on these two phenomena and to test the prediction that awareness would affect proprioceptive shift and adaptation in a correlated manner.

### Feature 2. Non-monotonic function of perceived hand position following introduction of visual perturbation

A core feature of PReMo is that the error signal is the difference between perceived hand position and desired hand position. Perceived hand position is rarely measured, perhaps because this variable is not relevant in visuo-centric models. When it is measured (e.g., in studies measuring proprioceptive shift), the data are usually obtained outside the context of adaptation (Cressman & Henriques, 2010). That is, these proprioceptive assays are taken before and after the block of adaptation trials, providing limited insight into the dynamics of this key component of PReMo: the perceived hand location *during* implicit adaptation.

We conducted a study to probe the time course of perceived hand position in a continuous manner. Participants reached to a target and received 15° clamped visual feedback. They were asked to maintain their terminal hand position after every reach and provide a verbal report of the angular position of their hand (Figure 7; note that this verbal report indexes the participant’s perceived hand position, 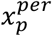; this measure is not the same as proprioceptive shift, *β*_*p*_) (Tsay, Parvin, et al., 2020) [Footnote 5: The original intent of the experiment was to directly test participant’s perceived hand position during adaptation, seeking to confirm the common assumption that participants are unaware of the effects of adaptation. The results of the study, especially the non-monotonic shape of the hand report function, inspired the development of PReMo.]. Surprisingly, these reports followed a striking, non-monotonic pattern. The initial responses were biased towards the clamped visual feedback by ∼5°, but then reversed direction, gradually shifting *away* from the clamped visual feedback and eventually plateauing at around 2° on the opposite side of the actual target position (Figure 7B).

**Figure 7.**
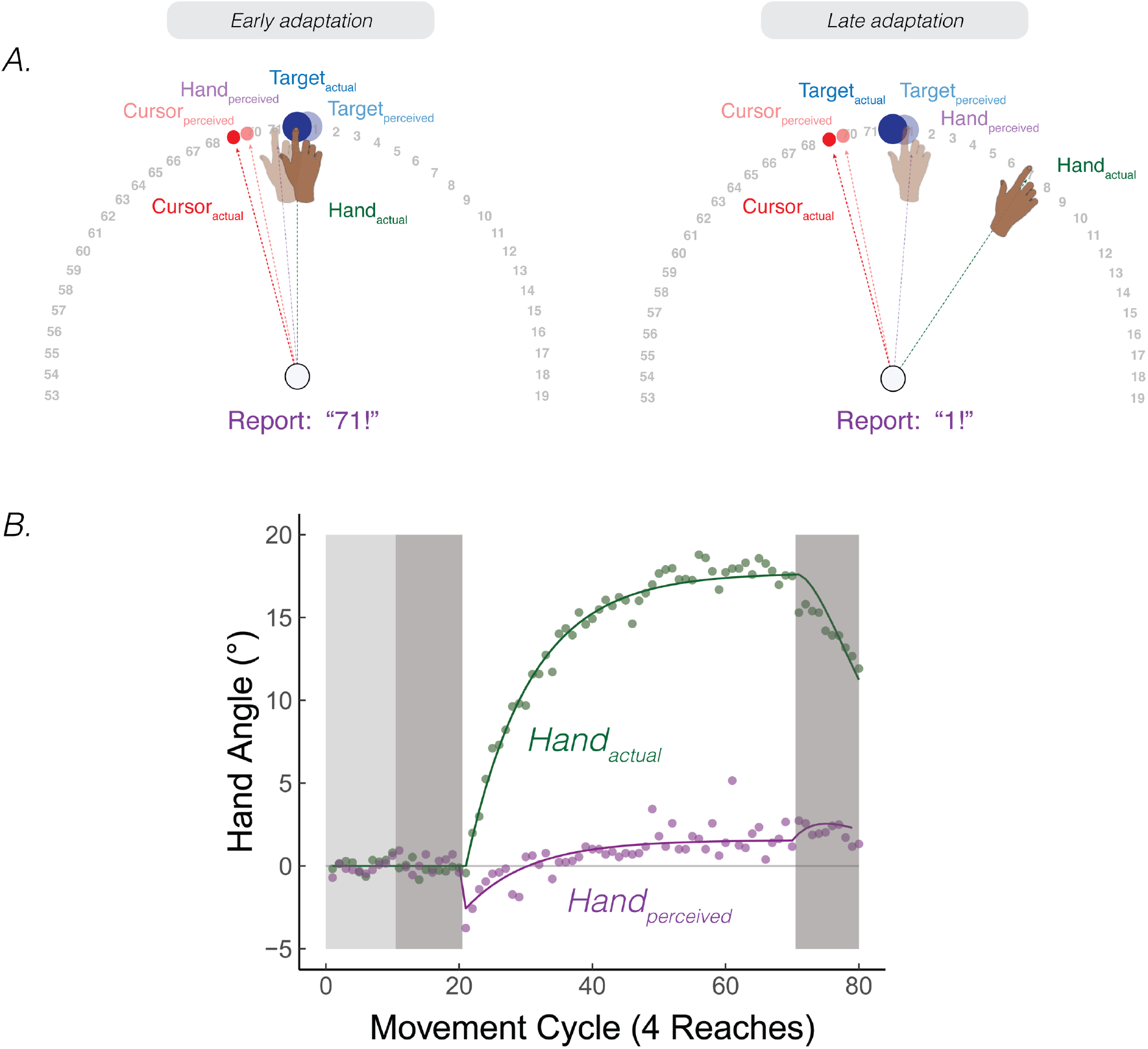
Continuous reports of perceived hand position during implicit adaptation. **A)** On each trial, participants reached to a target (blue dot) with and received 15° clamped visual feedback (red dot). After each reach, a number wheel appeared on the screen, prompting the participant to verbally report the angular position of their (unseen) hand. Participants perceived their hand on the left side of the target, shifted towards the clamped visual cursor early in adaptation. The left side shows the state of each variable contributing to perceived hand position right after the introduction of the clamp and the right side shows their states late in the adaptation block. **B)** After baseline trials (light grey = veridical feedback, dark grey = no feedback), participants exhibited 20° of implicit adaptation (green) while the hand reports (purple) showed an initial bias towards the visual cursor, followed by a reversal, eventually overshooting the target by ∼2°. Lines denote model fits of the proprioceptive re-alignment model (*R*^2^ = 0.995).

The shape and dynamics of this function are readily accounted for by PReMo. The initial shift towards the clamp is consistent with the size and rapid time course of the proprioceptive shift (Ruttle et al., 2021, 2018). It is this shift, a consequence of crossmodal recalibration between vision and proprioception, which introduces the proprioceptive error signal that drives adaptation (Eq 5 & 7). This error signal results in the hand moving in the opposite direction of the clamp in order to counter the perceived proprioceptive error. This, in turn, will result in a corresponding change in the perceived hand position since it is determined in part by the actual hand position (Eq 1 & 3). As such, the perceived hand location gradually converges with the perceived target location.

Intriguingly, the perceived hand position does not asymptote at the actual target location, but rather overshoots the target location by ∼1°. This overshoot of the perceived hand position is also accounted for by PReMo. During exposure to the rotated visual feedback, not only is the perceived location of the cursor shifted away from the actual cursor position due to cross-sensory recalibration, but this shift is assumed to be a generic shift in visual space, not specific to just the cursor (Simani et al., 2007) (see Feature 1, Corollary 2 above). As such, the perceived location of the target is shifted away from the cursor (Figure 7A-B). Indeed, the overshoot provides a measure of the magnitude of this visual shift, given the assumption that behavior will asymptote when the proprioceptive error is nullified (i.e., when the perceived hand position corresponds to the desired hand position: the perceived target location).

Notably, the putative 1° of visual shift inferred from the PReMo parameter fits is consistent with empirical estimates of a visual shift induced by a visuo-proprioceptive discrepancy (Rand & Heuer, 2019a; Simani et al., 2007). The convergence between measures obtained from very different experimental tasks supports a surprising feature of PReMo, namely that the final perceived hand position will be displaced from the actual target location. That being said, a more direct test would be to modify the continuous report task, asking participants to report the (remembered) location of the target rather than the hand.

### Feature 3. The effect of proprioceptive uncertainty on implicit adaptation

The preceding sections focused on how a proprioceptive shift biases the perceived hand position away from the movement goal, eliciting a proprioceptive error that drives implicit adaptation. Another factor influencing implicit adaptation is the variability in perceived hand location, i.e., proprioceptive uncertainty. This is also estimated from the psychometric function obtained from subjective reports of sensed hand position. If obtained during adaptation, the shift in perceived hand position would impact the estimates of variability. As such, a cleaner approach is to measure proprioceptive uncertainty prior to adaptation (Figure 3). As shown in several experiments, greater proprioceptive uncertainty/variability is associated with a greater extent of implicit adaptation (Figure 3H, I, J).

As with measures of proprioceptive shift, conventional visuo-centric models of implicit adaptation do not account for the relationship between proprioceptive uncertainty and the magnitude of adaptation. In these models, the extent of implicit adaptation reflects the point of equilibrium between learning and forgetting from a *visual* error and do not specify how the extent of implicit adaptation may be related to proprioception. In contrast, PReMo predicts that proprioceptive variability will be negatively correlated with implicit adaptation (Eq. 9). When there is greater uncertainty in the proprioceptive system, the perceived hand position is more biased by the location of sensory expectations from the motor command (i.e., the visual target, Eq 1). Therefore, participants with greater uncertainty in proprioception would require a greater change in their actual hand position to bring their perceived hand position into alignment with the perceived target.

### Feature 4. The effect of visual uncertainty on implicit adaptation

Visual uncertainty can also affect implicit adaptation. In a seminal study by Burge et al (2008), the visual feedback in a 6° (small) visuomotor rotation task was provided in the form of a sharply defined cursor (low uncertainty) or a diffuse Gaussian blob (high uncertainty). In the high uncertainty condition, implicit adaptation was attenuated both in rate and asymptotic value. The authors interpreted this effect through the lens of optimal integration (Burge et al., 2008; Ernst & Banks, 2002), where the learning rate is determined by the participant’s confidence in their estimate of the sensory prediction and feedback. When confidence in either is low, the learning rate will be decreased; thus, the added uncertainty introduced by the Gaussian blob reduces the learning rate and, consequently, the asymptotic value of final learning. By this view, visual uncertainty should attenuate implicit adaptation for all visual error sizes.

PReMo offers an alternative interpretation of these results. Rather than assume that visual uncertainty impacts the strength of the error signal, PReMo postulates that visual uncertainty indirectly affects implicit adaptation by influencing the magnitude of the proprioceptive shift. This hypothesis predicts that the impact of visual uncertainty may depend on the visual error size.

To explain this prediction, consider Eq 9, the core equation specifying the relationship between the upper bound of implicit adaptation and the degree of the proprioceptive shift, *β*_*p*_. When the visual error is small, the proprioceptive shift, being a fraction of the integrated hand/cursor positions will be below the level where it saturates (Eq 5). As such, we can substitute *β*_*p*_ with the expression, 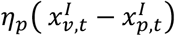, that is, a fraction of the difference between integrated positions of the hand and cursor (Eq 14):

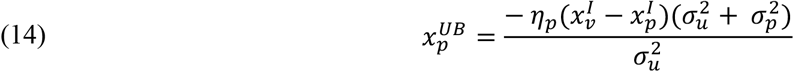

Furthermore, we can substitute the integrated positions of the hand 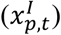 and the visual cursor 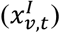 from Eq 1 and Eq 2, respectively, to relate the upper bound of implicit adaptation with uncertainty in proprioception 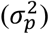, vision 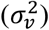, and the sensory prediction 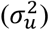. For the sake of simple exposition, we assumed that visual shifts are negligible and that participants continue to aim directly to the target (*G*_*t*_= 0), but the same logic would apply if visual shifts were non-zero (Eq 15):

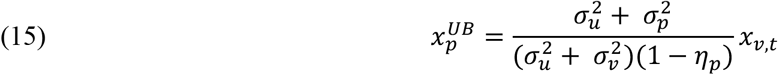

As visual uncertainty increases, the denominator in Eq 15 increases, and thus, the upper bound of implicit adaptation decreases. More specifically, when visual uncertainty of a small visual error increases, the integrated cursor is drawn closer to the visual target (the aiming location), and, thus, closer to the integrated hand position (assumed to be near the target during early adaptation). For small errors, the discrepancy between integrated positions of the cursor and hand decreases as visual uncertainty increases. This will reduce the size of the proprioceptive shift and, consequently, result in the attenuation of implicit adaptation.

In contrast, consider the situation when the visual error is large. Now the visuo-proprioceptive discrepancy is large 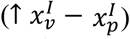 and we can assume the proprioceptive shift will be at the point of saturation (*β*_*p,sat*_). As such, the upper bound of implicit adaptation will no longer depend on visual uncertainty (see Eq 9) and, thus, implicit adaptation will not be attenuated by visual uncertainty. In summary, PReMo, predicts an interaction between error size and the effect of visual uncertainty on adaptation.

The results of an experiment in which we varied visual uncertainty and error size are consistent with this prediction. To have full control over the size of the error, we used the clamped feedback method. We varied visual uncertainty (cursor = certain feedback, Gaussian cloud = uncertain feedback) and the size of the visual error (3.5° = small error, 30° = large error) in a 2×2 design (Figure 8A). Visual uncertainty attenuated implicit adaptation when the error size was small (3.5°), convergent with the results of Burge et al (2008) (Figure 8B). However, visual uncertainty did not attenuate implicit adaptation when the error size was large (30°) (Tsay, Avraham, et al., 2020), yielding the predicted interaction.

**Figure 8.**
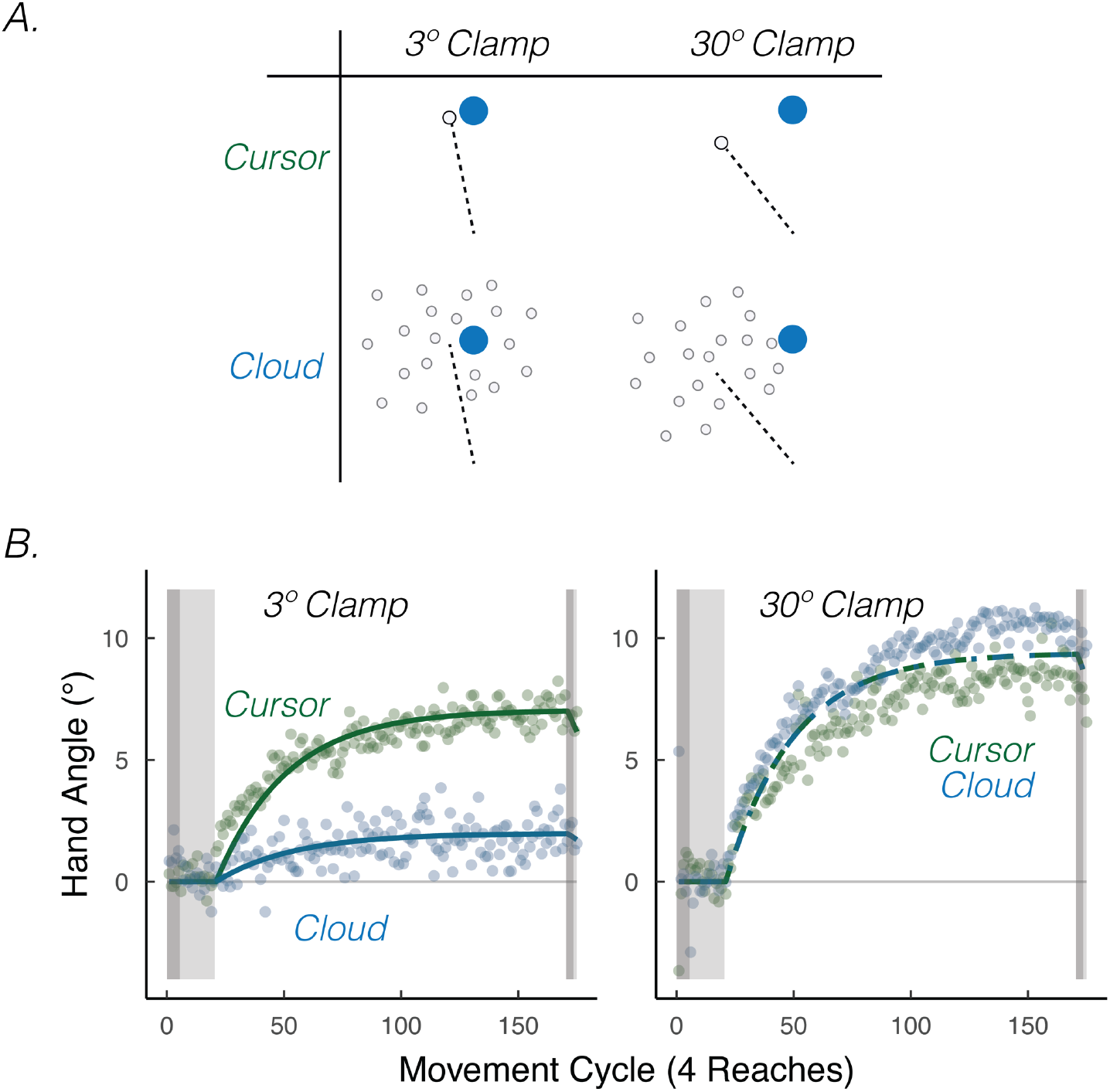
Visual uncertainty attenuates implicit motor adaptation in response to small visual errors, but not large visual errors. **A)** Experimental design in Tsay et al (2020). Participants made reaching movements in a similar setup as Figure 1A. Feedback was provided as a small 3.5° visual clamp or large 30° visual clamp, either in the form of a cursor or cloud (2×2 between-subject factorial design). **B)** Tsay et al (2020) Results. Implicit adaptation was attenuated by the cloud feedback when the clamp size was 3.5° but not when the clamp size was 30°. Lines denote model fits of the proprioceptive re-alignment model (*R*^2^ = 0.929).

It is possible that the Gaussian cloud led to lower adaptation because the added noise induced more “successful” trials, given previous work showing that implicit adaptation is attenuated when the visual cursor intersects the target (Leow et al., 2018, 2020; Tsay, Haith, et al., 2022). This concern was one of the reasons why we blanked the target at reach onset in Tsay et al 2020. Nonetheless, more direct evidence that the attenuation stems from uncertainty (and not reward) comes from an unpublished study involving participants with low vision and matched controls, allowing a test of the effect of uncertainty when it comes from an intrinsic source (the individual) rather than manipulating an extrinsic source (the Gaussian cloud) (Tsay, Tan, et al., 2022). The results exhibit the same interaction, with visual uncertainty due to low vision attenuating implicit adaptation for small errors but not large errors (despite visual feedback being equated in both groups).

### Feature 5. The effect of sensory prediction uncertainty on implicit adaptation

All models of implicit adaptation require a comparison of predicted and observed feedback to derive an error signal. In the previous section, we discussed how PReMo can account for attenuated adaptation observed in the face of noisy feedback – a variable that is easy to manipulate. In this section we consider the effects of noisy predictions, a latent measure in the model, and one that is difficult to manipulate.

*Extrinsic* and *intrinsic* sources of variability have been posited to impact the strength of the predicted sensory consequences of a movement. Extrinsic variability, defined here as variability in movement outcomes that are not attributable to one’s own motor output, will reduce one’s ability to make accurate sensory predictions. Such effects are usually simulated in the lab by varying the perturbation across trials. For example, Albert et al (2021) compared adaptation in two groups of participants, one exposed to a constant 30° visuomotor rotation and a second exposed to a variable rotation that, across trials, averaged 30° (SD = 12°). Adaptation was attenuated by around 30% in the latter condition. From the perspective of PReMo, this effect could be attributed to increased sensory prediction noise. However, these results should be interpreted with caution: Other studies have found no effect of perturbation variability on adaptation (Avraham, Keizman, et al., 2020; Butcher et al., 2017) or even an amplified effect on learning from increased perturbation variability (Burge et al., 2008). Moreover, the attenuation of adaptation due to uncertainly can emerge from differential sampling of error space relative to a condition with low uncertainty, even when the learning rate is identical for the two conditions (Wang et al., in preparation). Given the mixed results on this issue (also see: (Hutter & Taylor, 2018)), it is unclear if extrinsic variability contributes to the strength of the sensory prediction.

In contrast to extrinsic variability, we define intrinsic variability as noise arising within the agent’s nervous system. High intrinsic variability will, over trials, decrease the accuracy of the sensory predictions. Indeed, an impairment in generating a sensory prediction provides one mechanistic account of why individuals with cerebellar pathology show attenuated sensorimotor adaptation across a range of tasks (Donchin et al., 2012; Fernandez-Ruiz et al., 2007; Gibo et al., 2013; Hadjiosif et al., 2014; Izawa et al., 2012; Martin et al., 1996; Parrell et al., 2021; Schlerf et al., 2013; Tseng et al., 2007). From the perspective of a state-space model, this impairment is manifest as a lower learning rate (Figure 9), although this term encompasses a number of processes. PReMo suggests a specific interpretation: Noisier sensory predictions will result in the actual hand position having a relatively larger contribution to the perceived location of their hand. As such, a smaller change in actual hand position would be required to nullify the proprioceptive error (Eqs 9, 10), effectively lowering the upper bound of implicit adaptation.

**Figure 9.**
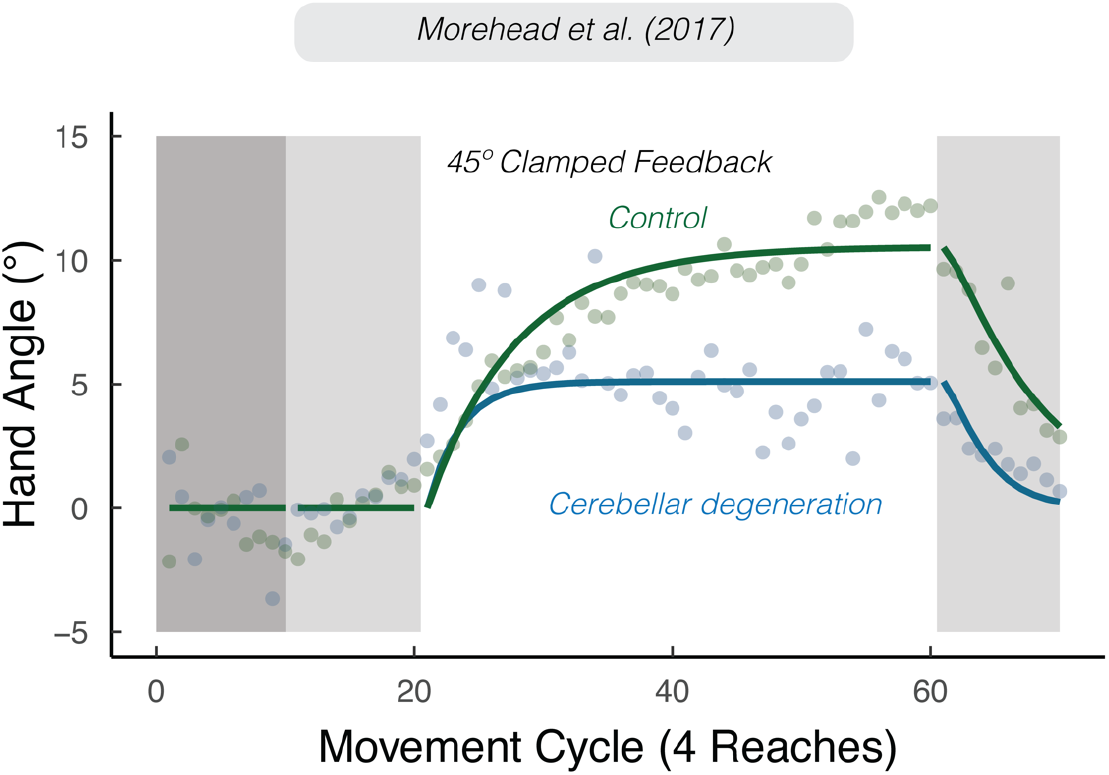
Sensory prediction uncertainty attenuates implicit adaptation. Attenuated adaptation in individuals with cerebellar degeneration compared to matched controls in response to 45° clamped feedback. Data from Morehead et al (2017). Lines denote model fits of the proprioceptive re-alignment model (*R*^2^ = 0.897).

Evidence from a number of different tasks is consistent with the hypothesis that cerebellar pathology is associated with noisier sensory predictions (Bhanpuri et al., 2013; Gaffin-Cahn et al., 2019; Therrien & Bastian, 2019; Weeks et al., 2017b). Nonetheless, we recognize that cerebellar pathology may disrupt other processes relevant for implicit adaptation. For example, the disease process may result in a lower (generic) learning rate or core proprioceptive variables. However, with respect to the latter, various lines of evidence indicate that proprioception, at least those aspects highlighted in PReMo, are not impacted by cerebellar pathology. First, these individuals do not exhibit impairment on measures of proprioception obtained without volitional movement (i.e., under static conditions) (Bhanpuri et al., 2013). Second, the magnitude of the proprioceptive shift is comparable to that observed in control participants (Henriques et al., 2014). Third, proprioceptive variability appears to be comparable between individuals with cerebellar pathology and matched controls (Bhanpuri et al., 2013; Weeks et al., 2017a, 2017b).

### Feature 6: Generalizing the proprioceptive re-alignment model from visuomotor to force-field adaptation

Implicit adaptation is observed over a wide range of contexts, reflecting the importance of keeping the sensorimotor system precisely calibrated. In terms of arm movements, force-field perturbations have provided a second model task to study adaptation (Shadmehr et al., 1993). In a typical task, participants reach to a visual target while holding the handle of a robotic device. The robot is programmed such that it exerts a velocity-dependent force in a direction orthogonal to the hand’s movement. Over the course of learning, participants come to exert an opposing time-varying force, resulting in a trajectory that once again follows a relatively straight path to the target.

In contrast to visuomotor adaptation tasks, there is no manipulation of sensory feedback in a typical force-field study; people see and feel their hand exactly where it is throughout the experiment. Nevertheless, several studies have reported sensory shifts following force-field adaptation (Figure 10A). In particular, the perceived hand position becomes shifted in the direction of the force-field (Mattar et al., 2013; Ohashi, Gribble, et al., 2019; Ostry et al., 2010). [Footnote 6: At odds with this pattern, Haith et al. (Haith et al., 2009) reported a proprioceptive shift during force-field adaptation but in the direction opposite to the applied force. However, this study only involved a leftward force-field. The rightward shifts in perceived hand position may be due to a systematic rightward proprioceptive drift, a phenomenon observed in right-handed participants with repeated reaches, with or without feedback (Brown et al., 2003a, 2003b).].

**Figure 10.**
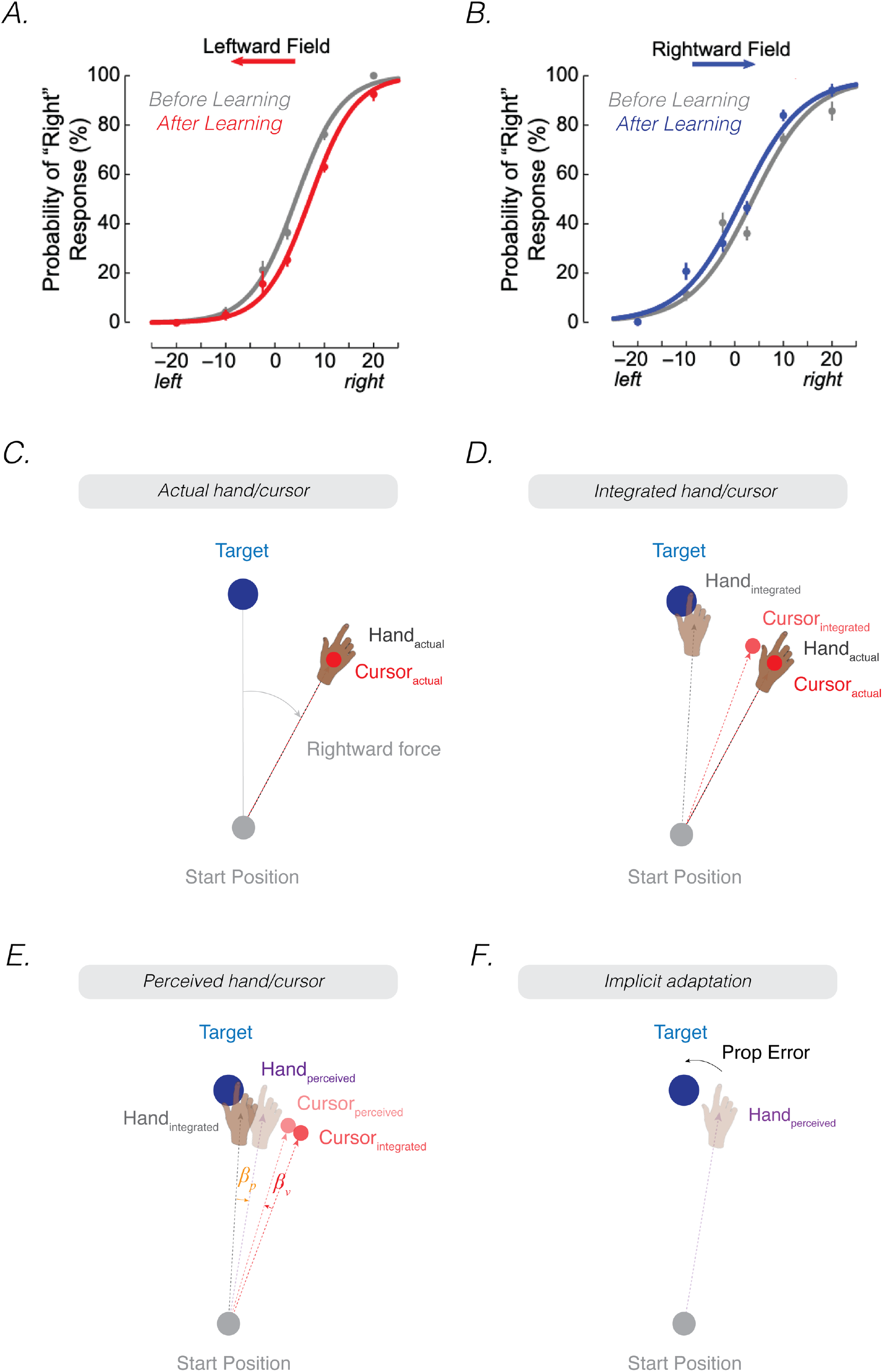
The proprioceptive re-alignment model explains sensory and motor changes during force-field adaptation. **A, B)** Participants’ perceptual judgments of actual hand position are biased in the direction of the recently experienced force-field (Ostry et al, 2010). These psychometric curves were obtained using a staircase method performed before and after force-field adaptation: the participant made center-out reaching movements toward a target. Force channels pushed the participant’s hand towards the left or right of the target by varying amounts. At the end of the movement, the participant judged the position of their hand relative to the target (left or right). Each participant’s shift quantified as the change in the point of subjective equality (PSE). The shift in the PSE in panels A and B indicate that the perceived hand position following force-field adaptation was shifted in the direction of the force-field. **C)** Upon introduction of the force-field perturbation, the hand and (veridical) cursor are displaced in the direction of the force-field, especially at peak velocity. Note that the hand and cursor positions are illustrated at the endpoint position for ease of exposition. **D)** Assuming that proprioceptive uncertainty is greater than visual uncertainty, the integrated hand position would be closer to the target than the integrated cursor position due to principles of optimal integration. **E)** The two integrated positions then mutually calibrate, resulting in a proprioceptive shift (*β*_*p*_) of the integrated hand towards the integrated cursor position, and a visual shift (*β*_*v*_)of the integrated cursor position towards the integrated hand position, forming the perceived hand and perceived cursor locations. **F)** The proprioceptive error (mismatch between the perceived hand position and the target) drives adaptation, a force profile in the opposition direction of the force-field.

PReMo can account for the shift in perceived hand position in the direction of the force-field. Consider the situation where a force-field pushes the hand to the right, an effect that is maximal at peak velocity for a velocity dependent force-field (Figures 10C-F). An estimate of hand position will involve intramodal integration of sensory feedback signals (the cursor for vision, the actual hand position for proprioception) and the sensory prediction (i.e., a straight trajectory of the hand towards the target). This will result in an integrated representation of the hand and cursor trajectory as shifted towards the target. Importantly, given that proprioceptive variability is greater than visual variability (van Beers et al., 2002; Van Beers et al., 1999), this shift towards the target is likely greater for the hand than the cursor. The visuo-proprioceptive discrepancy between integrated positions will result in crossmodal calibration, with the integrated hand and cursor positions shifted towards each other, forming the perceived hand and cursor positions, respectively (Figure 10E). Thus, there will be a small but systematic shift in the perceived location of the hand in the direction of the force-field perturbation.

Models of force-field adaptation suggest that the error signal driving adaptation is the deviation between the ideal and actual forces applied during the movement. This can be estimated based on the deviation of the hand’s trajectory from a straight line (Donchin et al., 2003). These models are agnostic to whether this error is fundamentally visual or proprioceptive since the position of the cursor and hand are one and the same. That is, the error could be visual (the trajectory of the cursor was not straight towards the target) or proprioceptive (the trajectory of the hand did not feel straight towards the target). Interestingly, neurologically healthy and congenitally blind individuals can adapt to a force-field perturbation without the aid of vision, relying solely on proprioceptive input (DiZio & Lackner, 2000). Furthermore, when opposing visual and proprioceptive errors are provided, aftereffects measured during the no-feedback block after adaptation are in the direction counteracting the proprioceptive error instead of the visual error (Hayashi et al., 2020). As such, we suggest that force-field adaptation may be fundamentally proprioceptive. Consistent with the basic premise of PReMo, the difference between the perceived and desired hand position constitutes the error signal to drive force-field adaptation, a process that can operate in the absence of visual feedback. Strikingly, Matter et al (2013) found that the degree in which participants adapt to the forcefield is correlated with the amount of proprioceptive shift in the direction of the force-field, strengthening the link between the sensory and motor changes that arise during force-field adaptation (also see Feature 1).

In summary, PReMo offers a unified account of the motor and perceptual changes observed during force-field and visuomotor adaptation – both of which place emphasis on participants reaching directly to a target. The applicability of the model for other types of movements remains to be seen. Proprioception seems quite relevant for locomotor adaptation where the goal of the motor system is to maintain gait symmetry (Morton & Bastian, 2006; Reisman et al., 2007; Rossi et al., 2019): The misalignment between the desired and perceived gait might serve as a proprioceptive error, triggering implicit locomotive adaptation to restore its symmetry. Indeed, for locomotor adaptation, it is unclear what sort of visual information might be used to derive an error signal. In contrast, the goal in saccade adaptation is fundamentally visual, to align the eye on a target (Groh & Sparks, 1996; Grüsser, 1983; Pélisson et al., 2010). The oculomotor system appears to rely on a visual error signal to maintain calibration (Lewis et al., 2001; Noto & Robinson, 2001; Wallman & Fuchs, 1998).

## VII. Concluding remarks

In the current article, we have proposed a model in which proprioception is the key driver of implicit adaptation. In contrast to the current visuo-centric zeitgeist, we have argued that adaptation can be best understood as minimizing a proprioceptive error, the discrepancy between the perceived limb position and its intended goal. On ecological grounds, our model reframes adaptation in terms of the primary intention of most manual actions, namely, to use our hands to interact and manipulate objects in the world. In visuo-centric models, the central goal is achieved in an indirect manner, with the error signal derived from visual feedback about the movement outcome being the primary agent of change. Empirically, the proprioceptive re-alignment model accounts for a wide range of unexplained, and in some cases, unintuitive phenomena: Changes in proprioception observed during both visuomotor and force-field adaptation, phenomenal experience of perceived hand position, the effect of goal and sensory uncertainty on adaptation, and saturation effects observed in the rate and extent of implicit adaptation.

To be clear, the core ideas of PReMo are framed at Marr’s ‘computational’ and ‘algorithmic’ levels of explanation. At the computational level, we seek to explain *why* implicit adaptation is elicited (to align felt hand position with the movement goal); at the algorithmic level, we ask *how* implicit adaptation is instantiated (felt hand position being a combination of vision, proprioception of the moving limb, and efferent information; movement goal being the perceived location of the target). We hope PReMo motivates studies that focus on the implementational level. Here we anticipate that it will be important to consider both peripheral (Dimitriou, 2016) and central mechanisms (Latash, 2021; Proske & Gandevia, 2012) to account for the modification of multisensory representations across the course of adaptation.

It should be emphasized that, to date, much of the key evidence for PReMo comes from correlational studies; in particular, the relationship between the magnitude of adaptation and the extent and variability of induced changes in proprioception following adaptation. While studies using a wide range of methods have revealed robust correlations between measures of proprioception and adaptation, other studies have failed to find significant correlations (Cressman et al., 2021, 2010) (Vandevoorde & Orban de Xivry, 2021). The reason for these differences is unclear but may be related to the methodological differences. For instance, when proprioception is assessed via subjective reports obtained after an active movement, it may be impossible to dissociate the relative contributions of proprioceptive variability and sensory prediction variability to perceived hand position (Izawa et al., 2012; Izawa & Shadmehr, 2011; Synofzik et al., 2008). More important, correlations of the proprioceptive data with implicit adaptation would confound these two sources of variability.

Development and validation of proprioceptive measures that do not rely on subjective reports may bypass these shortcomings. Rand and Heuer, for instance, have developed a measure of proprioception that is based on movement kinematics, without participants being aware of the assessment (Rand & Heuer, 2019b). Using a center-out reaching task, the participants’ perceived hand position after the outbound movement is inferred by the angular trajectory of the inbound movement back to the start position. A straight trajectory to the start position may indicate that the participant is fully aware of their hand position, whereas any deviation in this trajectory is inferred to reveal a proprioceptive bias. This indirect measure of proprioception shows the signature of a proprioceptive shift with a similar time course as that observed when the shift is measured in a more direct manner.

Even though each correlation, when considered in isolation, should be interpreted with caution, we believe that when considered in aggregate, PReMo provides a parsimonious explanation for a wide range of empirical phenomena. Importantly, PReMo provides sufficient detail to generate a host of qualitative and quantitative predictions based on a reasonable set of assumptions. For example, the computation of perceived hand position is based on established principles of sensory integration (Ernst & Banks, 2002) and sensory recalibration (Zaidel et al., 2011). Moreover, the key proposition of PReMo, namely that implicit sensorimotor adaptation operates to reduce a proprioceptive error, is highly ecological.

As with all correlational work, inferences about the causality are indirect and may be obscured by mediating variables. For example, instead of implicit adaptation being driven by the proprioceptive shift, it is possible that the visual error introduced by a perturbation independently drives implicit adaptation and the proprioceptive shift. By laying out a broad range of phenomena in this review, we hope to establish a benchmark for comparing the relative merits of PReMo and alternative hypotheses.

Beyond model comparison and appeals to parsimony, a more direct tack to evaluate the core proposition of PReMo would involve experimental manipulations of proprioception. Brain stimulation methods have been used to perturb central mechanisms for proprioception (Armenta Salas et al., 2018; Balslev et al., 2004, 2007; Block et al., 2013; Miall et al., 2007) and tendon vibration has been a fruitful way to perturb proprioceptive signals arising from the periphery (Bernier et al., 2007; Gilhodes et al., 1986; Goodwin et al., 1972; Manzone & Tremblay, 2020; Roll et al., 1991). PReMo would predict that implicit adaptation would be enhanced with greater proprioceptive bias induced by tendon vibration to one muscle group (e.g., vibration to the biceps resulting in illusory elbow extension) and greater proprioceptive uncertainty via vibrating opposing muscle groups (e.g., vibration to biceps and triceps adding noise to peripheral proprioceptive afferents).

Adaptation encompasses a critical feature of our motor competence, the ability to use our hands to interact and manipulate the environment. As experimentalists we introduce non-ecological perturbations to probe the system, with the principles that emerge from these studies shedding insight into those processes essential for maintaining a precisely calibrated sensorimotor system. This process operates in an obligatory and rigid manner, responding, according to our model, to the mismatch between the desired and perceived proprioceptive feedback. As noted throughout this review, the extent of this recalibration process is limited, likely reflecting the natural statistics of proprioceptive errors. The model does not capture the full range of motor capabilities we exhibit as humans (Listman et al., 2021). Skill learning requires a much more flexible system, one that can exploit multiple sources of information and heuristics to create novel movement patterns (Yang et al., 2021). These capabilities draw on multiple learning processes that use a broad range of error and reinforcement signals (Galea et al., 2011; Shmuelof et al., 2012; Tsay, Kim, Saxena, et al., 2021) that may be attuned to different contexts (Avraham, Taylor, et al., 2020; Heald et al., 2021). While we anticipate that these processes are sensitive to multimodal inputs, it will be useful to revisit these models with an eye on the relevance of proprioception.

**Open Questions**

1. How general are the principles of PReMo for understanding sensorimotor adaptation in other motor domains? For example, can PReMo account for locomotor adaptation?
2. How do we reconcile PReMo’s emphasis on proprioceptive error with the observation that individuals with severe proprioceptive deficits exhibit adaptation? Is this due to a compensatory process? Or the internal representation of hand position based on sensory expectancies interacting with biases from vision? Insights into this question will also be relevant for recalibrating movements when learning to use a tool, prosthetic limb, or body-augmentation devices in which the goal of the action is not isomorphic with a proprioceptive signal.
3. A core principle of PReMo is that the perceived location of the target and hand are biased by various sources of information. What is the impact of these biases on other learning processes engaged during sensorimotor adaptation tasks (e.g., use-dependent learning or strategic re-aiming)?
4. We have proposed that the impairment in adaptation associated with cerebellar pathology may arise from noisier sensory predictions, a hypothesis consistent with the view that the cerebellum is essential for predicting the proprioceptive outcome of a movement based on efference copy. Alternatively, within the framework of PReMo, the impairment might relate to a reduced learning rate, a disturbance of proprioception, or a combination of factors. Specifying the source of impairment will require experiments involving tasks that yield independent measures of these variables to constrain parameters when fitting learning functions.
5. New insights into cerebellar function have come about by considering the representation of error signals in Purkinje cells during saccade adaptation (Herzfeld et al., 2018). Can the principles of PReMo be validated neurophysiologically by examining the time course of error-related activity during adaptation. For example, in response to clamped feedback, PReMo would predict an attenuation of the error signal as proprioceptive alignment occurs whereas standard state-space models would predict little change, with the asymptote reached when the effect of the persistent error is offset by forgetting.
6. What are the implications of PReMo for physical rehabilitation of neurologic populations who are at high risk of proprioceptive impairments, such as stroke and Parkinson’s disease?

## Appendix

### The standard visuo-centric model

Sensorimotor adaptation is often expressed as a state-space model, positing that implicit adaptation is a process of trial-to-trial learning from a visual error as well as trial-to-trial forgetting. The learning process is controlled by a learning rate (*e*), which specifies how much is learned from the visual error (*K*). The forgetting process is controlled by a retention parameter (*A*). These two processes dictate how the participant’s state (*x*) (e.g., hand trajectory) changes over time, from trial t to trial t + 1.

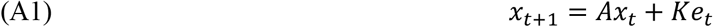

The upper bound of implicit adaptation (*x*_*UB*_) is achieved when *x*_*t +1*_ is equal to *x*_*t*_.

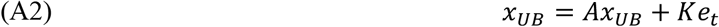

By rearranging these terms, one can appreciate that this upper bound is achieved when the amount of forgetting (1 - A) is equal to the amount of learning from the visual error:

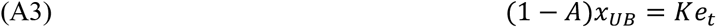

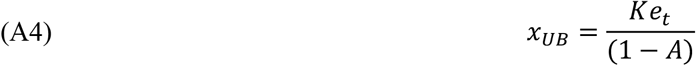

### Fitting the Proprioceptive re-alignment model

Using R language’s fmincon function, we started with 10 different initial sets of parameter values to estimate the parameter values that minimized the least squared error between the average data and model output. The key dependent variables were hand angle and reports of perceived hand position. The eight key parameters were *σ*_*v*_ (visual uncertainty), *σ*_*p*_ (proprioceptive uncertainty), *σ*_*u*_ (sensory prediction/expectation uncertainty), *㭲*_*p*_ (proprioceptive shift ratio), *σ*_*v*_ (visual shift ratio), *β*_*p,sat*_ (saturation of proprioceptive shift), *K* (learning rate), and *A* (rate of proprioceptive decay).

### Proprioceptive shift does not correlate with proprioceptive variability

While it is reasonable to posit that proprioceptive shift and proprioceptive variability will be correlated with one another given that each variable correlates with the extent of implicit adaptation, this need not be the case: Two variables can be independent from one another, yet still both correlate with a third variable. For example, shift and uncertainty could each make positive, yet independent contributions to implicit adaptation. There are theoretical reasons, mainly from the sensory integration world to expect shift and uncertainty to be correlated. Empirically, however, several studies have found this to not be the case (Ayala et al., 2020; Tsay, Kim, Parvin, et al., 2021). While noting this is a null result, these data suggest that the degree of sensory recalibration may not follow a Bayesian optimal rule in which the extent of sensory shifts are based on the relative reliabilities of each sensory signal. This hypothesis is consistent with Zaidel et al (2011) who found that visual/vestibular cross-modal recalibration did not follow Bayesian optimality principles. Instead, these two sensory modalities appear to shift towards each other in a fixed ratio manner. For these reasons, we opted to formulate the proprioceptive shift in PReMo as independent of sensory uncertainty (see Eqs 5 & 6).

## Acknowledgements

We thank members of the CognAc lab, Sensorimotor learning lab, and BLAM lab for insightful discussions. We also thank Amanda Therrien, Romeo Chua, and Cristina Rossi for their constructive feedback on the manuscript.

